# Optimization of closed-loop electrical stimulation enables robust cerebellar-directed seizure control

**DOI:** 10.1101/2021.09.03.458908

**Authors:** Bethany J. Stieve, Thomas J. Richner, Chris Krook-Magnuson, Theoden I. Netoff, Esther Krook-Magnuson

**Affiliations:** Graduate Program in Neuroscience, University of Minnesota, Minneapolis 55455, United States; Department of Neurology, Mayo Clinic, Rochester 55902, United States; Department of Biomedical Engineering, University of Minnesota, Minneapolis 55455, United States; Department of Neuroscience, University of Minnesota, Minneapolis 55455, United States

**Keywords:** personalized medicine, responsive neurostimulation, search algorithm, DBS, neuromodulation

## Abstract

Additional treatment options for temporal lobe epilepsy are needed, and potential interventions targeting the cerebellum are of interest. Previous animal work has shown strong inhibition of hippocampal seizures through on-demand optogenetic manipulation of the cerebellum. However, decades of work examining electrical stimulation – a more immediately translatable approach – targeting the cerebellum has produced very mixed results. We were therefore interested in exploring the impact that stimulation parameters may have on seizure outcomes. Using a mouse model of temporal lobe epilepsy, we conducted on-demand electrical stimulation of the cerebellar cortex, and varied stimulation charge, frequency, and pulse width, resulting in over a thousand different potential combinations of settings. To explore this parameter space in an efficient, data-driven, manner, we utilized Bayesian optimization with Gaussian process regression, implemented in Matlab with an Expected Improvement Plus acquisition function. We examined two different fitting conditions and two different electrode orientations. Following the optimization process, we conducted additional on-demand experiments to test the effectiveness of selected settings. Across all animals, we found that Bayesian optimization allowed identification of effective intervention settings. Additionally, generally similar optimal settings were identified across animals, suggesting that personalized optimization may not always be necessary. While optimal settings were consistently effective, stimulation with settings predicted from the Gaussian process regression to be ineffective failed to provide seizure control. Taken together, our results provide a blueprint for exploration of a large parameter space for seizure control, and illustrate that robust inhibition of seizures *can* be achieved with electrical stimulation of the cerebellum, but only if the correct stimulation parameters are used.

## Introduction

Temporal lobe epilepsy, the most common form of epilepsy in adults^1^, is notoriously pharmacoresistent, leading the majority of patients to have a severe decline in their quality of life^2,3^. Less than half of epilepsy patients experience seizure control with antiseizure medications^4^, leading many to seek neurostimulation adjunct therapies, including deep brain stimulation (DBS), responsive neurostimulation (RNS), and vagal nerve stimulation (VNS). However, even with these additional therapeutic options, many patients continue to have uncontrolled seizures. Illustrating patient need, in the initial RNS cohort, the average seizure frequency prior to initiating RNS treatment was over 50 seizures a month, despite on-going pharmacological approaches, one third of patients having been treated with VNS, and one third of patients already having had neurosurgery^5^. Unfortunately, while RNS was beneficial to many of those patients, less than 20% experienced even one year of seizure freedom over a nine-year period^5^. Clearly, additional therapy options are needed.

The cerebellum is classically considered a motor structure^6^, but is increasingly appreciated for its role in a range of functions^7-11^. The cerebellum has been a potential intervention target of interest for epilepsy for decades^12-16^. Recently, on-demand optogenetic work has suggested robust inhibition of both hippocampal^17-19^ and absence seizures^20,21^ in rodent models with cerebellar directed intervention. However, electrical stimulation of the cerebellum has produced very mixed results, both in human clinical studies^22-33^ and in various animal models^34-54^. On-demand optogenetic approaches have a number of experimental benefits over open-loop electrical stimulation, including cell-type and temporal specificity, as well as direct control over the direction of modulation of neurons (excitation versus inhibition). However, optogenetic methods are not currently a clinical option for epilepsy patients (although exciting progress is being made on making this a potential future reality^55^). Therefore, we were interested in determining if electrical stimulation of the cerebellum could be done in a manner which provided robust inhibition of seizures.

When conducting electrical stimulation interventions, a range of parameters must be decided, often with insufficient *a priori* knowledge to guide the selection. For example, electrical stimulation can vary the frequency of stimulation, the strength of the stimulation, and the duration of each pulse of stimulation (among other variables). We hypothesized that these settings may be critical in determining outcomes of electrical stimulation of the cerebellum for seizure control. This includes the potential for interactions between variables (e.g. frequency and pulse width). Considering the combinatorial nature, the number of potential settings to test quickly becomes overwhelming.

We therefore turned to Bayesian optimization^56-58^. Bayesian optimization allows for a thorough, data driven, exploration of large parameter spaces, and is being increasingly applied to neuroscience related areas, including epilepsy^59,60^ and DBS^61-64^. We used a mouse model of temporal lobe epilepsy, detected spontaneous seizures online via hippocampal local field potential (LFP) recordings, and delivered on-demand electrical stimulation to the cerebellar vermis. We varied the frequency, charge, and pulse width, and computed effectiveness (measured as seizure duration) online. In parallel, we completed Gaussian process regression of the incoming data, allowing selection of the next set of parameters to try based on the modelling of the effectiveness across the parameter space. The next set of parameters tested balanced exploration of areas predicted to be effective, and sufficient exploration of the full parameter space, so as to avoid mistaking local minima for a global minimum, and to achieve a more complete picture of the parameter space. After identifying parameters predicted to be effective (optimized settings), we completed additional testing to directly compare seizure outcomes with this optimized intervention to no intervention. For each mouse we additionally tested a set of parameters predicted to be ineffective. We tested different initial fitting conditions, and different electrode orientations. Across our experiments, we found that robust seizure inhibition could be achieved, but only with effective stimulation settings. Across animals, modeling conditions, and electrode orientations, interventions using high frequency stimulation and relatively strong charge were identified as optimal and were able to provide robust seizure inhibition. Our findings illustrate the benefits of taking a Bayesian optimization approach, and help resolve controversies around electrical stimulation of the cerebellum for seizure control. We show that electrical stimulation of the cerebellum can be highly effective, but outcomes are critically dependent on the stimulation parameters.

## Materials and methods

### Animals

For all experiments male and female C57BL/6J (Jackson Laboratories stock #000664) mice, bred in-house, were used. Mice were sexed at the time of weaning based on external genitalia. Prior to electrographic recordings, mice lived in standard housing conditions in the animal facility at the University of Minnesota. Following kainate (KA) injections, male mice were singly housed to avoid fighting. After electrode implantation, all mice were singly housed to avoid damage to implants. During electrographic recordings, mice lived in investigator-managed housing. At all times, mice had *ad libitum* access to food and water, and were on a 12 h light: 12 h dark (/low red light) cycle. All experimental protocols were approved by the University of Minnesota’s Institutional Animal Care and Use Committee (IACUC protocol 2011-238662A).

### Stereotactic surgeries

#### Epilepsy induction

The mouse unilateral intra-hippocampal KA model of chronic temporal lobe epilepsy^65,66^ was implemented^67^. In brief, mice (postnatal day 45 or later) were anesthetized with isoflurane and received local bupivacaine. Stereotaxic injection of 100 nL of 18.5 mM KA in sterile saline into the hippocampus (from bregma, AP: −2.0 mm, ML 1.25 mm left, DV: 1.6 mm) was delivered with a microliter syringe (Hamilton 2 μl Neuros, Reno, NV). Mice were rapidly recovered from anesthesia (within 5 minutes of injection)^68^ and treated with NeoPredef. As expected, spontaneous recurrent electrographic seizures were observed within several weeks post-injection^65^. Optimization experiments started on average 12.1 ± 2.7 weeks post-KA injection (range: 3.9 – 57.3 weeks).

#### Electrode implantation

After at least 12 days post KA-injection, mice were implanted with two electrodes: a bipolar twisted microwire electrode (50 µm diameter stainless steel with polyimide insulation, P1 Technologies, Roanoke, VA) for depth electrographic recordings in the hippocampus, and a low-impedance bipolar stimulation electrode (200 µm diameter tungsten with polyimide insulation, P1 Technologies) on the cerebellum. Prior to implantation, stimulating electrodes were fashioned by bending a pair of untwisted electrodes 90 degrees and trimming the ends to 0.5 mm beyond the bend, creating electrode feet 1 mm apart. The polyimide insulation along the 0.5 mm end was exposed with a blade. Under isoflurane anesthesia with carprofen (5 mg/kg subcutaneous) and local subcutaneous bupivacaine, mice were implanted with the recording electrode in the hippocampus (from bregma, AP: −2.6 mm, ML 1.75 mm left, DV: 1.4 mm) and the stimulating electrode epidurally along the midline, over cerebellar lobules 4/5 and 6. For the stimulating electrode, a 2.5 mm craniotomy, 6.75 mm posterior to bregma, along the midline was drilled. The electrode was placed in the center of the craniotomy, with the feet positioned either perpendicular or parallel to the midline, lightly pressed into the dura and then sealed over with UV curing dental acrylic (Pentron FlowIt, Brea, CA), insulating the dorsal surface of the electrodes, but keeping the ventral surface exposed to the tissue. Both recording and stimulating electrodes were secured with Metabond acrylic. Mice were treated with Neopredef and analgesics (ibuprofen, 50-80 mg/kg/day dissolved in drinking water, for 1 pre-surgical and 3 post-surgical days), and allowed to recover for at least a week prior to electrophysiology recordings.

### Real-time Bayesian optimization of on-demand seizure-intervention

#### Electrophysiological recordings and electrical stimulation

Epileptiform events were detected in real-time and used to trigger trains of current pulses applied to the cerebellar cortex (see online seizure detection below). Mice were tethered and freely behaving in a modified home cage during optimization and testing. Hippocampal LFPs were recorded using a BrownLee 440 amplifier (settings: 1-2000 Hz passband, 1 MOhm input impedance, 300-fold gain). These settings were selected to ensure stimulation artifacts never saturated input of the National Instruments data acquisition device (USB-6229 BNC). The analog inputs and outputs were sampled and generated at 20 kHz with 16-bit resolution so that the artifacts could be recorded and removed, preserving the LFPs with no gaps (see below and **Supplemental Figure 1**). Pulse trains were generated on-demand by a digital to analog converter (DAQ, National Instruments, 6229 (BNC), Austin, TX) and converted into current-controlled pulses at 100 µA/V by an A-M Systems 2200 stimulator (A-M Systems, Sequim, WA). To avoid any non-zero voltage drift at the output, a first-order RC high pass filter (either R = 0.5 MOhm, C = 215 nF, fc = 1.5 Hz; or R = 1MOhm, C = 215 nF, fc = 0.75 Hz) was installed at the output of the stimulator. To ensure multi-day battery life, the stimulator’s batteries were replaced with large sealed lead acid batteries (12V, 7AH).

#### Online seizure detection

LFPs were processed in real-time to detect and record seizure durations with custom Matlab (Mathworks, Natick, MA) based software, using a series of digital filters, rectification, and thresholding to isolate epileptic spikes (**Supplemental Figure 1**). Ictal spike detection was individualized for each mouse based on several features, including minimum and maximum inter-spike distance, spike width, and spike amplitude (similar to Armstrong et al.^67^). Additionally, epileptic spikes were often associated with brief high frequency oscillations (HFOs)^69^; therefore, in some instances, an HFO detector (500-1500 Hz; using the same features as spike detection; spike width, amplitude, etc.) was used to improve spike discrimination. Spike and HFO coincidence detection was individualized, as well as the minimum number of spikes arriving in two seconds to constitute the beginning -- or for online seizure duration analysis (see below) the end -- of an event. In order to observe spikes during stimulation, the stimulus-locked artifacts were removed from the LFPs using a least mean squares adaptive filter^70^ with 20 ms of history at a 20 kHz sampling rate (Matlab’s DSP toolbox, **Supplemental Figure 1**). The adaptive filter fits a finite impulse response (FIR) filter by comparing the stimulus waveform to the recorded signal’s artifacts, similar to template removal, but generalized across waveforms. In order to enable the artifact-removal processing, and support optimization of up to four mice in parallel, data was processed in 1 s segments. Note that this caused a 1 s delay between seizure detection and delivery of stimulation. The performance and stability of the adaptive filter was improved by reapplying the adaptive filter three times to each second of data. The first two passes were used to adapt the filter and the artifact was removed in the third pass. In this manner, even artifacts near the beginning of the train were profoundly suppressed. Over fitting was not an issue; epileptiform spikes were not correlated in time to the stimulus onset and therefore did not appear to be affected by the adaptive filter (**Supplemental Figure 1**). For the optimization process, the duration of epileptiform events were defined as the time between the first and last epileptic spike of the event, and converted to a logarithmic scale using the natural log.

#### Subject-specific Gaussian process regression and Bayesian optimization

In real-time, we optimized cerebellar stimulation for suppression of hippocampal seizures. In brief, stimulation of the cerebellar cortex was delivered in response to seizure detection. The duration of the seizure was recorded and estimated using Gaussian process regression to form a response surface. The response surface predicts seizure duration as a function of stimulation parameters, across the entire parameter space, and can make estimates for even unsampled parameter combinations. Based on the response-surface, Bayesian optimization determines which parameters to test next, to efficiently explore the parameter space in a data-driven manner^61,71^ (**Fig. 1A**).

**Figure 1.**
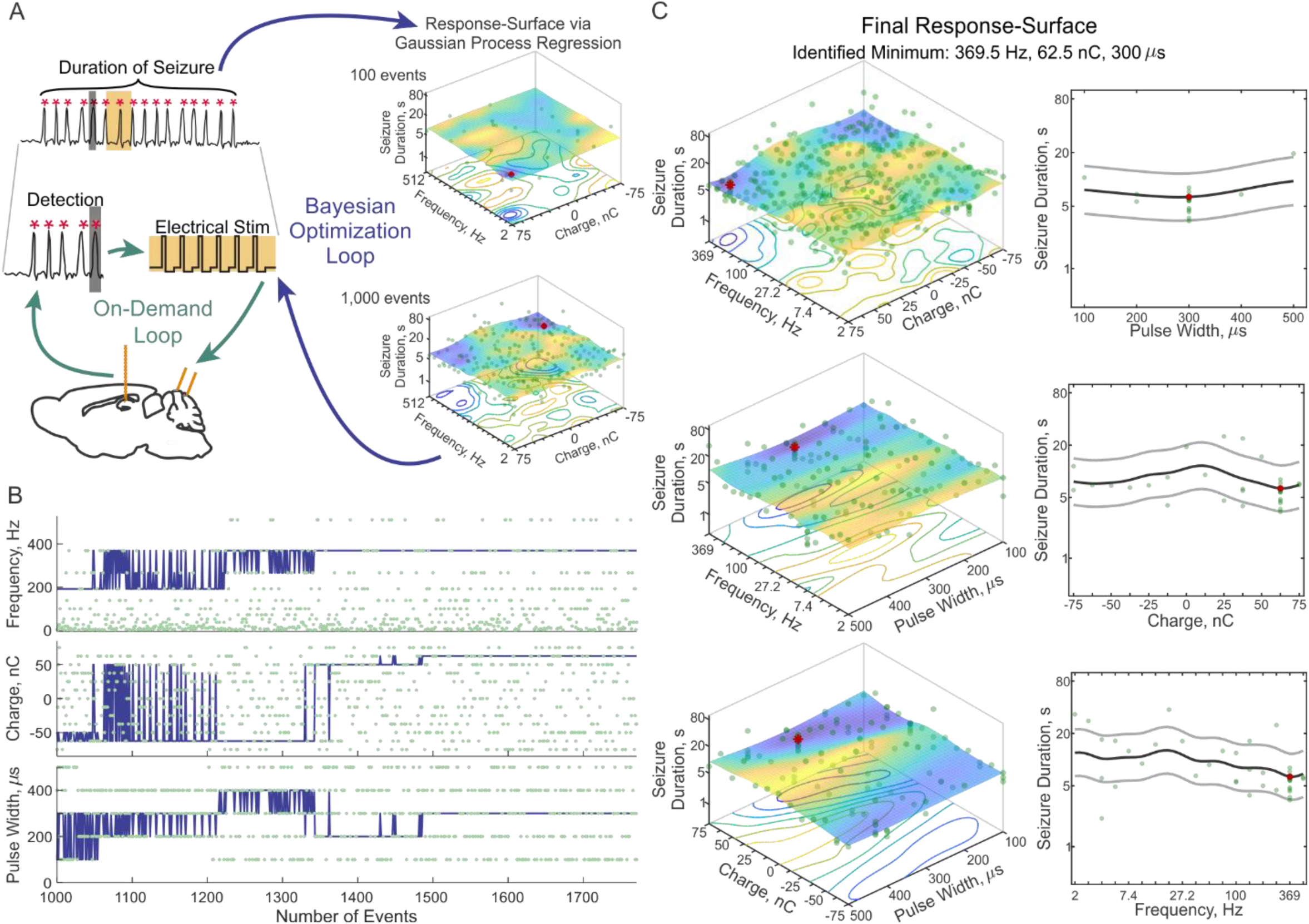
On-demand Bayesian optimization of cerebellar-directed seizure-intervention. (**A**) Online hippocampal seizure detection triggers electrical stimulation of the cerebellar cortex (on-demand loop). Seizure-duration is recorded in real-time and Gaussian process regression is used to build a response-surface (Bayesian optimization loop). The response-surface is updated with each event, and the next set of parameters to test is determined via Bayesian optimization. Note that only frequency and charge are illustrated here, but optimization occurred over frequency, charge, and pulse width. (**B**) Example parameter walk demonstrating that the optimizer selects a range of parameters (green dots), and updates the estimate of the best set of parameters (blue line) with each event. In this example, the optimizer settles on 369.5 Hz, 62.5 nC, and 300 µs, but continues to test several other combinations of settings. (**C**) Final response-surface of the same example animal as in (*B*), where cooler and warmer colors represent shorter and longer event durations, respectively. The red dot indicates the identified minimum, representing the optimal parameter. Due to the high dimensionality of the data, three planes are shown. In the top row, the interaction of frequency versus charge for combinations with 300 µs pulse width is shown on the left, while the impact of pulse width, at 369.5 Hz and 62.5 nC, is shown on the right. The next rows follow the same pattern, highlighting the interaction of frequency and pulse width, and the impact of charge (*middle row*), and the interaction of pulse width and charge, and the impact of frequency (*bottom row*). Note that in subsequent figures, only one plane for a given response-surface is illustrated, through the identified minimum.

In further detail, we optimized across three stimulation parameters for seizure-suppression, using two different parameter spacing and fitting conditions. The optimized parameters included *i*) frequency (2 to 512 Hz, 18 or 9 steps spaced logarithmically), *ii*) charge per pulse (−75 to 75 nC, 13 or 7 steps spaced linearly), and *iii*) pulse width (100-500 µs, 5 steps spaced linearly). In addition to non-stimulation (zero charge) combinations, the discretized parameter space contained either 1,080 or 270 stimulation (non-zero) parameter combinations. Stimulation parameters that were kept constant included the current waveform (biphasic), stimulation train duration (3 seconds; used in previous optogenetic experiments^17-19^), and the pulse asymmetry amplitude phase ratio (3:1), as illustrated in **Supplemental Figure 1**. The charge range was chosen to remain within the safety guidelines reviewed by Cogan et al.^72^ with an estimated electrode area of 170,000 µm^2^ which is between that of a micro- and macroelectrode. Charge was balanced in each biphasic pulse to avoid adverse electrochemical reactions^72^.

Bayesian optimization on a Gaussian process regression is a sequential method for exploring a space to find an optimum in as few steps as possible^56-58^. Gaussian process regression, like the more familiar linear regression, fits a surface to data points, but unlike a linear regression, Gaussian process regression is allowed to bend locally to better fit the data. The flexibility of the response-surface is determined by the length-scale of the kernel function. Smaller length-scales fit the observed data more tightly, reducing the influence of neighboring observations on expected outcomes of parameters, resulting in a more flexible response-surface. Larger length-scales allow for greater influence of previous observations, resulting in a more generalized, smoother surface. To start to determine whether effective optimization would depend on the length-scale, we changed the length-scale with the number of steps in the frequency and charge dimensions. In both parameter spaces, a Matern 5/2 kernel was used, and pulse width (5 steps) had a length-scale of 150 µs. In the parameter space with 1,080 non-zero combinations (18 steps for frequency, 13 steps for charge), length-scales were 3.4 dB-Hz for the frequency and 15 nC for charge. In the parameter space with 270 non-zero combinations (9 steps for frequency, 7 steps for charge), length-scales increased to 7.22 dB-Hz for the frequency and 30 nC for charge, resulting in a relatively less flexible parameter space (more generalized), compared to the first parameter space.

At every point in space, the Gaussian process regression estimates the mean and confidence of the optimized variable. Bayesian optimization reads the response-surface created by the Gaussian process regression to decide what setting to test next. The optimizer must balance testing in areas that were previously found to be successful to improve confidence and find incremental improvement (reducing time spent in ineffective areas), and exploring more widely for a potentially better setting elsewhere in the parameter space (to prevent missing the global minimum). This tradeoff is essential to the nature of optimization problems. To balance this tradeoff, we used the Expected-Improvement Plus acquisition function (EI-plus; Matlab’s BayesianOptimization class^73^). EI-plus chooses settings expected to most greatly improve the optimum, with an additional comparison to avoid over-exploitation of a local minimum. Using a term called the exploration ratio (set at 0.5 for our study), the optimizer determines, based on the variance of the noise and the confidence of the estimate, whether the selected point in the parameter space is being over-exploited, and if so, modifies the kernel function and selects a new point^74^. Progress of optimization was periodically monitored, and typically considered stable if the optimizer chose the same set of the three parameters for at least 100 consecutive events, after a minimum of 1,000 events was collected. If over 1,500 events were collected, and the optimizer had not stabilized at the time of observation, the last parameter combination was used as the optimized set. Optimization took on average 5.5 ± 0.45 days (range: 3 – 9 days).

#### Testing of optimized settings

Following optimization, we conducted a second round of testing using a single-loop on-demand approach (i.e., only the “on-demand loop” in **Fig. 1A**) to obtain a sufficient number of events for direct statistical comparisons. For each seizure detected, either *i*) optimized stimulation, *ii*) non-optimal stimulation, or *iii*) no stimulation (as an internal control) was delivered in an interleaved, random fashion. The minimum of each animal’s individualized response-surface, associated with shortest seizure durations, was used as the optimized parameter set. The maximum of the final response-surface, the point associated with the longest seizure durations, was used as the non-optimal parameter set. Note that for one animal (parallel electrode orientation, more flexible parameter space), the non-optimal setting was 0 nC, and therefore the same as the no-stimulation control (this animal is represented by a dark orange square in the inset in figure 4). In all other cases a non-zero charge was identified as the non-optimal setting.

Given that response-surfaces followed similar patterns across animals, we conducted an additional round of testing with the perpendicular electrode orientation in the more flexible parameter space, to test if group-derived settings would provide sufficient seizure-suppression, compared to individualized settings. Four conditions were tested with the same, randomly interleaved, on-demand approach: *i*) the individualized optimized setting for each mouse, *ii*) the group mode setting, *iii*) the group average setting, and *iv*) no-stimulation. The group mode setting was based on the most common setting in each of the three parameters (**Table 1**). Given that charge values spanned both sides of zero, and surfaces suggested mirroring, to calculate the mode optimal charge, first, the absolute value of charge was used. Then, to determine the polarity, the more common sign was selected. If two values were equally common, one was arbitrarily chosen to serve as the mode. The group average setting was derived from the minimum of the average response-surface (**Fig. 2F**). Average response-surfaces were calculated by taking the average of normalized final response-surfaces from all mice with the same electrode orientation, within the same parameter space, with each animal weighted equally.

**Table 1.** Summary of data for individual animals presented in Figure 2.

| Animal ID, Sex<sup>a</sup> | Optimized Parameters |  |  |  |  | Non-Optimal Parameters |  |  |  |  |
| --- | --- | --- | --- | --- | --- | --- | --- | --- | --- | --- |
|  | Freq. (Hz) | Charge (nC) | Pulse Width (μs) | ΔSeizure Duration (%)<sup>b</sup> | p-values (KS; MW)<sup>c</sup> | Freq. (Hz) | Charge (nC) | Pulse Width (μs) | ΔSeizure Duration (%) | p-values (KS; MW) |
| Yellow ◆<br>♀ | 266.7 | -75 | 400 | -82 | <u>7.9E-47;</u><br><u>8.9E-37</u> | 27.2 | -12.5 | 100 | 5 | 0.46;<br>0.97 |
| D. Orange ■ ♂ | 512 | -62.5 | 100 | -92 | <u>8.9E-50;</u><br><u>1.3E-43</u> | 3.8 | 12.5 | 400 | -5 | 0.84;<br>0.91 |
| Green ▼<br>♂ | 369.5 | 62.5 | 300 | -87 | <u>3.0E-73;</u><br><u>5.8E-61</u> | 138.9 | -25 | 400 | 4 | 0.43;<br>0.89 |
| L. Orange ●<br>♂ | 512 | 50 | 400 | -71 | <u>3.6E-36;</u><br><u>1.8E-37</u> | 2.8 | 50 | 500 | -12 | 0.13;<br>0.06 |
| D. Blue ◀<br>♂ | 266.7 | -62.5 | 300 | -76 | <u>1.4E-28;</u><br><u>4.6E-22</u> | 14.2 | -12.5 | 100 | 8 | 0.67;<br>0.46 |
| L. Blue ▲<br>♀ | 266.7 | 75 | 200 | -28 | <u>1.5E-08;</u><br><u>3.1E-06</u> | 10.2 | -62.5 | 200 | -6 | 0.62;<br>0.54 |
| Pink ▶<br>♂ | 192.4 | -75 | 400 | -59 | <u>3.9E-46;</u><br><u>1.1E-33</u> | 7.4 | -25 | 300 | 1 | 0.40;<br>0.56 |

<sup>a</sup>Animal sex is denoted by ♂ for male, ♀ for female. Color and shape indicate how the animal is presented in Figure 2 (L=light, D=dark).
<sup>b</sup>ΔSeizure Duration is calculated relative to the average post-stimulation-onset seizure duration of the no stimulation control.
<sup>c</sup>Significant values (p-value < 0.05) are underlined in the table.

**Figure 2.**
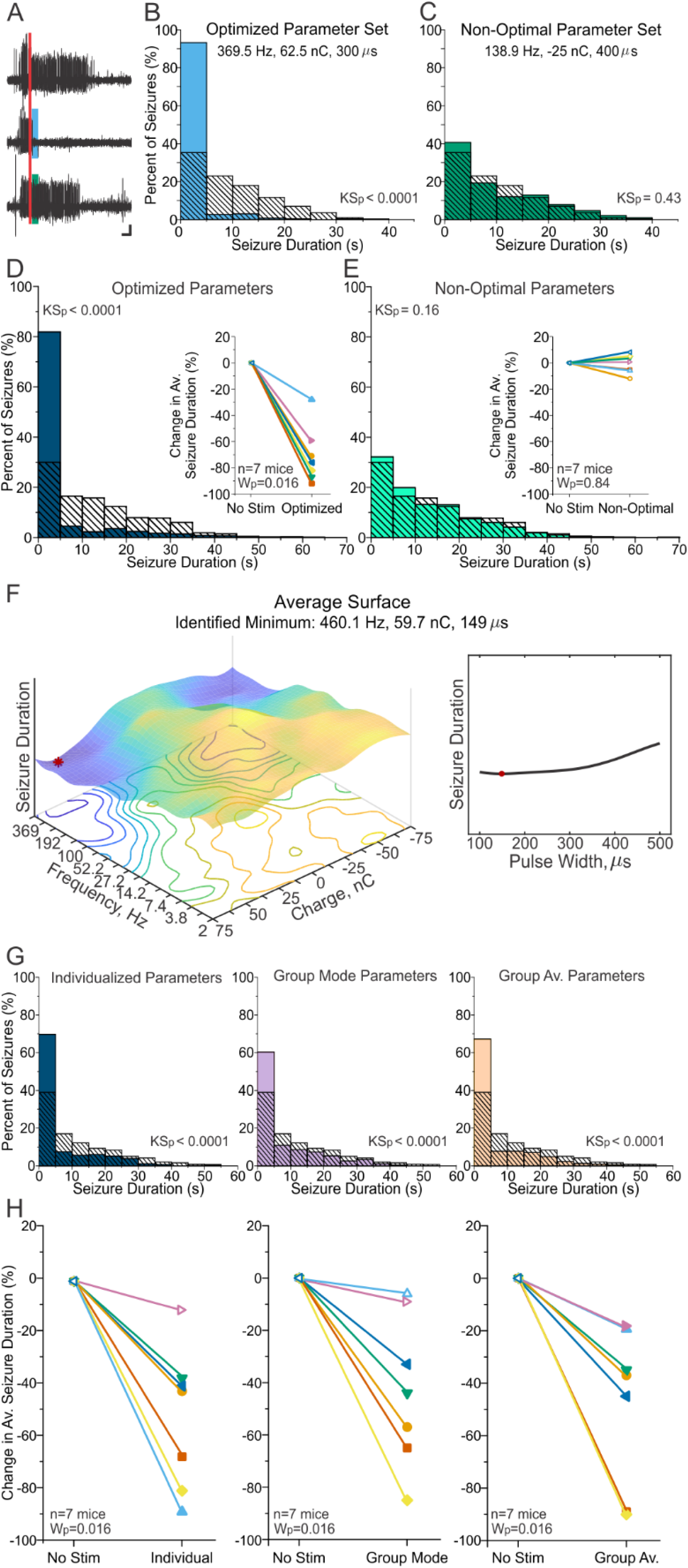
Electrical stimulation of the cerebellum with optimized parameters provides seizure-control. (**A**) Example seizure events from the same animal shown in Figure 1. Events were detected online (red bar) and received, in a random, interleaved fashion, no stimulation (*top*), stimulation with optimal parameters (*middle*, blue bar), or stimulation with non-optimal parameters (*bottom*, green bar). Stimulation was delivered for three seconds, with frequency, charge, and pulse width determined by the final response-surface (see Fig. 1C). Scale bar: 5 s, 0.5 mV. (**B** and **C**) Post-stimulation-onset seizure duration distributions for the example animal shown in (*A*). (**B**) Seizure duration distributions when stimulation was applied with the optimized parameter set of 369.5 Hz frequency, 62.5 nC charge, and 300 µs pulse width (light blue bars) versus no-stimulation (hashed bars). In this example animal, an 87% reduction in seizure duration occurred (*p* < 0.0001, KS test). (**C**) Seizure duration distributions for the non-optimal parameter set (dark green bars; in this animal: 138.9 Hz, −25 nC, and 400 µs pulse width), or for comparison, no-stimulation (hashed bars). In this animal, a non-significant 4% increase in seizure duration was noted with non-optimal stimulation (p = 0.43, KS test). Note that the same no-stimulation seizure duration distribution is overlaid in (*B*) and (*C*). (**D** and **E**) Post-stimulation-onset seizure duration distributions across animals (n=7 mice), using 100 randomly chosen seizure events per condition per animal, and with the same no-stimulation data set illustrated for comparison (hashed bars) in both (*D*) and (*E*). (**D**) Dark blue bars: events receiving each animal’s optimized stimulation; hashed bars: no-stimulation control. Stimulation with optimized parameters provides significant and consistent seizure-reduction (inset: significant at the individual level in 7 of 7 animals tested; average 70.7 ± 8.2% reduction in seizure duration, p = 0.016, Wilcoxon). (**E**) Light green bars: events with non-optimal parameters; hashed bars: no stimulation control. Inset: No significant change in seizure duration with non-optimal parameters (insignificant change in 7 of 7 animals; on average a 0.7 ± 2.7% reduction in seizure duration, *p* = 0.84, Wilcoxon). (**F**) Average response-surface, based on the final response-surfaces of the 7 mice. Red dot marks the minimum of the surface at 460.1 Hz, 59.7 nC, and 149 µs pulse width. (**G**) Group-level post-stimulation-onset seizure duration distributions; 100 random seizure events per condition per animal. Dark blue bars: Seizure duration when stimulation was applied with each animal’s individualized optimized parameter set. Purple bars: Seizure duration with stimulation settings based on the most commonly identified optimal parameters (group mode parameters; 266.7 Hz, - 62.5 nC, 400 µs pulse width; see Table 1). Orange bars: Seizure duration with stimulation settings based on the minimum of the average surface (460.1 Hz, 59.7 nC charge, 149 us pulse width, see (*F*)). Hashed bars: no stimulation control. (**H**) Stimulation provided with optimized parameters significantly reduces seizure duration whether parameters are derived from individualized optimization (*left*, average 52.1±10.3% reduction, p = 0.016 Wilcoxon), or group-derived settings based on the mode (*middle*, average 42.7±10.0% reduction; *p* = 0.016 Wilcoxon) or the minimum of the average surface (*right*, 47.4±11.4 % reduction, *p* = 0.016 Wilcoxon). At the group level, there was no significant difference in seizure-reduction between any of the three optimized stimulation conditions (p>0.375 for each comparison, Wilcoxons). Note that each color/symbol represents a different animal, with color coding consistent throughout the panel. Open symbols: insignificant change at the individual animal level (p>0.05). Closed symbols: significant change (p<0.01) for KS and MW tests at the individual animal level.

### Statistical analyses

During Bayesian optimization and single-loop data collection sessions, seizure duration was calculated on-line. Automated seizure-analysis was carefully monitored by manual inspection, and for single-loop data sets, results were confirmed through offline seizure duration analysis. Offline seizure duration analysis was completed using a combination of manual and automated methods, as described previously in Streng and Krook-Magnuson^18^. The stimulation condition was hidden to the reviewer (blinded analysis). Seizure duration from the onset of stimulation (1 second following seizure detection) was used for all offline statistical analyses. Given typical interspike intervals for ictal spikes, seizures that stopped > 500 msec prior to the onset of stimulation were excluded, and durations of seizures that stopped within 500 msec of stimulation were considered equal to 0 s. A minimum of 100 seizures per condition per animal were used for analysis. Previous power analyses indicate this provides a power of 0.82 for Mann-Whitney (MW) and 0.89 for two-sample Kolmogorov-Smirnov (KS) tests for a 20% decrease in seizure durations^17^.

Change in average seizure duration was calculated by subtracting the average no-stimulation duration from the stimulation duration, and dividing by the no-stimulation duration. A Friedman ANOVA was used to determine, at the group level, if there were any differences in seizure durations between conditions. Degrees of freedom were equal to 2 for Friedman ANOVAs comparing optimal, non-optimal, and no stimulation and 3 for Friedman ANOVA comparing personalized optimal, group mode settings, group average, and no stimulation. If the Freidman ANOVA resulted in a p-value < 0.05, then paired Wilcoxon tests were used for direct comparisons between conditions (note that only Wilcoxon values are presented). For group comparisons between non-paired data (e.g. electrode orientation), a Mann-Whitney was used. As an additional measure of group-level results, we sampled 100 events (randomly selected via Matlab) per condition per animal, and created summative histograms: post-stimulation-onset seizure durations were combined from each animal per stimulation condition, and compared to the associated no-stimulation distribution. Note that at the group-level the percentage of seizures stopping within 5s was calculated from this randomized subset of the data. Time from the end of a detected seizure until the start of the next seizure event was additionally calculated offline, and underwent statistical analysis in a similar manner as seizure duration (**Supplemental Fig. 3, Supplemental Tables 1, 3, 5**).

Statistical analyses were conducted using Matlab and Origin software. All statistical tests were conducted as two-tailed, and a p-value of < 0.05 was considered statistically significant. Results are reported as mean ± SEM.

### Data availability

Data presented in this manuscript is available upon any reasonable request. A version of the software for online seizure detection and Bayesian optimization, as well as the software used for offline seizure duration analysis, is available through github (https://github.com/KM-Lab).

## Results

Motivated by the success of on-demand optogenetic approaches targeting the cerebellum^17-21^, we sought to determine if an on-demand electrical stimulation approach could be effective in terminating temporal lobe seizures. Given the previous mixed literature on electrical stimulation of the cerebellum for seizure control^12,15^, we hypothesized that on-demand electrical stimulation of the cerebellum could be consistently effective, but only with the correct combination of stimulation parameters. As previous work using on-demand optogenetics used the dorsal intrahippocampal KA mouse model of temporal lobe epilepsy^17-19^, we similarly utilized this model. Spontaneous seizure events in chronically epileptic animals were detected on-line from LFP recordings from the hippocampus, and on-demand electrical stimulation was delivered to the midline cerebellum (vermis lobules 4/5 and 6) (**Figure 1A**, “on-demand loop”). We first tested stimulation with the electrode feet perpendicular to the midline using biphasic current pulse trains, and examined three different stimulation parameters: 1) frequency, 2) charge per pulse, and 3) pulse width. Specifically, we examined 2 Hz to 512 Hz stimulation, with 18 logarithmically spaced steps, charge from −75 nC to +75 nC, with 13 linearly spaced steps, and pulse widths from 100 µs to 500 µs, in 5 linearly spaced steps; this parameter space contained 1,080 non-zero stimulation parameter combinations. Given this large number of possible combinations of stimulation settings, we utilized Gaussian process regression to fit response surfaces and Bayesian optimization (**Fig. 1A**, “Bayesian optimization loop”) to effectively search the parameter space in a data driven manner.

The implemented subject-specific, personalized, search of the parameter space was designed to maximize data collection near areas showing good seizure inhibition (i.e, currently identified minima in seizure durations; blue line in **Fig.1B** and red dot on illustrated surfaces) and areas for which relatively less information about effectiveness was available, using an expected-improvement plus acquisition function^73,74^, as outlined in the Methods section. This allowed both good coverage of the space and increased confidence of effectiveness around identified minima, while avoiding over-exploitation of local minima. An example of a parameter walk is illustrated in **Figure 1B**, showing both the sampled settings (green dots; note the range of tested settings), and the running estimate of the best set of parameters (blue line; note that in this example the program settles on 369.5Hz, 62.5nC, and 300µs, starting around 1,500 events, but continues to sample other combinations of settings).

Despite potential non-stationarity effects and the relatively high variability in seizure durations in this model of epilepsy, Bayesian optimization successfully identified stimulation parameters for effective seizure-control. The progression of the response-surface for an example animal is shown in **Supplementary Video 1**, and the same animal’s final surface is illustrated in **Figure 1C**, with cooler colors illustrating shorter event durations and the red dots indicating the identified minimum (i.e., the predicted optimal combination of settings); note that given the high dimensionality of the data, three planes are shown. The top left plot illustrates the interaction of frequency versus charge (for combinations with 300µs pulse duration), and top right shows the impact of pulse width (at 369.5Hz and 62.5nC, i.e. the identified best combination for this animal). The middle plots show the same animal’s data set, but instead plots the interaction of frequency with pulse width (at 62.5nC), with the impact of charge shown to the right. Finally, the bottom row illustrates the interaction of charge and pulse width (at 369.5Hz), with the impact of frequency shown to the right. From these graphs, it is apparent that stronger charges (of either polarity) and higher frequency stimulation tend to provide more robust seizure control, with pulse width tending to play a smaller role in modulating seizure duration.

Using this optimization protocol, we were able to identify, for each animal, a combination of frequency, charge, and pulse width settings that were predicted to provide seizure inhibition (**Supplemental Figure 2; Table 1**). To validate the effectiveness of the individualized, optimized, parameter set we completed an additional set of experiments with each animal, utilizing a more traditional testing framework. Specifically, seizure detection resulted in either no stimulation (as a control), electrical stimulation to the cerebellum using that animal’s personalized optimal combination of settings (‘optimized’), or, as an additional control, intervention using a combination of parameters which, based on the previous Gaussian process regression, was predicted to be ineffective (‘non-optimal’) (**Figure 2**). Specifically, non-optimal settings were associated with the longest seizure durations in the response-surface (e.g. the maxima). These were delivered in an interleaved, random, manner. This experimental setup provided a sufficient number of events with each of these three conditions to allow for direct comparison. The identified personalized best set of parameters provided strong seizure inhibition (70.7±8.2% decrease in post-stimulation-onset seizure duration vs no intervention; n=7 mice; p=0.016 Wilcoxon; 82% of seizures stopping within 5s of the start of intervention; **Fig. 2D; Table 1**), confirming that our Bayesian optimization approach successfully identified a set of parameters capable of providing seizure inhibition. In contrast, non-optimal stimulation settings failed to provide seizure control (0.7±2.7% decrease vs no intervention; n=7 mice; p=0.84; at the individual animal level, insignificant change in seizure duration in all 7 of 7 tested animals; **Fig. 2E; Table 1**). Accordingly, optimized settings produced significantly better seizure inhibition than non-optimized settings (percent seizure reduction optimized vs non-optimized: p=0.016, Wilcoxon). We additionally examined the time to the next seizure event^17^. We found that optimized stimulation did not produce a consistent effect on the average time to next seizure, but did change the overall distribution of time to next seizure (**Supplemental Figure 3; Supplementary Table 1**). Non-optimal stimulation typically had no effect on the time to next seizure (**Supplemental Figure 3**).

Together these results illustrate that electrical stimulation of the cerebellum can be an effective approach to seizure inhibition, and only if the correct combination of stimulation parameters is utilized.

### Potential for non-individualized cerebellar stimulation settings

Across the animals tested, we noted similar final response surfaces (**Supplemental Figure 2; Table 1; Fig. 2F**). This suggested that, at least for similar seizure etiologies, once a set of effective settings is identified, it may be applicable in a broader setting. To test this, we completed an additional round of post-hoc testing. In this set of experiments, upon seizure detection, animals received either *i*) no stimulation, *ii*)personalized optimized, where the parameters selected were the optimized parameters from the subject’s own data *iii*)group mode, where the parameters were the most common (mode) charge, frequency, and pulse width identified as being optimal (specifically, 266.7 Hz frequency, −62.5 nC charge, 400 µs pulse width; **Table 1**), or *iv*) group average, where the parameters were selected as the minimum of the averaged response surface (specifically, 460.1 Hz frequency, 59.7 nC charge, 149 µs pulse width; red dot in **Fig. 2F**; note these specific frequency, charge, and pulse width settings were not directly tested during optimization, and were derived entirely from the Gaussian process regression). An expanded view with three planes of the average response-surface is illustrated in **Supplemental Figure 4**.

As before, these stimulations (or no-stimulation controls), occurred in an on-demand manner (selectively at the time of seizures), in a random, interleaved, order. While some animals showed greater seizure control with one parameter set versus another (**Fig. 2H**), across the animals, all three sets of stimulation parameters (personalized optimal, group mode, group average) produced strong inhibition of seizures (individualized vs no intervention: 52.1±10.3% decrease, p=0.016 Wilcoxon, n=7 animals, 70% of seizures stopping within 5s of intervention; group mode vs no intervention: 42.7±10.0% decrease, p=0.016 Wilcoxon, n=7 animals, 60% of seizures stopping within 5s of intervention; group average vs no intervention: 47.4±11.4% decrease, p=0.016 Wilcoxon, n=7 animals, 67% of seizures stopping within 5s of intervention; **Fig.2 G-H**). At the group level, there was no significant difference in outcomes between personalized optimization, group mode optimal settings, and group average optimal settings (p>0.375 for each comparison, Wilcoxon’s). This further underscores that within a similar population, similar cerebellar stimulation settings can generally be effective, and therefore, once a good set of parameters is identified, personalized optimization may not always be critical.

### Optimization process robust to different parameter spacing and fitting conditions

In initializing these experiments, there were a number of decisions made which could potentially impact the findings. These include how many gradations to try (i.e. step size) and how much each outcome for a given set of settings would impact neighboring combinations (i.e. the length-scale). We were interested in determining if the optimization process was sufficiently robust to allow for some variation in these hyperparameters, or if it was particularly sensitive to the fitting conditions. We therefore completed an additional set of experiments, again using Gaussian process regression and Bayesian optimization, but now with larger step sizes and effective length-scales, essentially allowing for a less flexible (i.e. more generalized) surface.

We found that the process was robust to these changes, and while more smoothed surfaces were generated (as expected), effective minima were still identified (**Figure 3; Supplemental Figure 5**). Identified optimal settings again provided robust inhibition of seizures (78.7±4.4% decrease in seizure duration vs no intervention; n=6 mice; p=0.031 Wilcoxon; 88% of seizures stopping within 5s of the start of intervention; **Fig. 3D; Supplemental Tables 2-3**), while non-optimal settings failed to inhibit seizures (9.8±3.5% decrease in seizure duration vs no intervention; n=6 mice; p=0.063 Wilcoxon; 33% of seizures stopping within 5s of the start of intervention; **Fig. 3E; Supplemental Tables 2-3**). Surfaces between animals showed similar profiles (**Supplemental Figure 5**). On average, as seen with a more flexible surface, a fairly high frequency (group average: 125.2 Hz), a fairly strong charge (71.9 nC), and fairly short pulse width (149 µs) was determined to be most effective (**Fig. 3F**).

**Figure 3.**
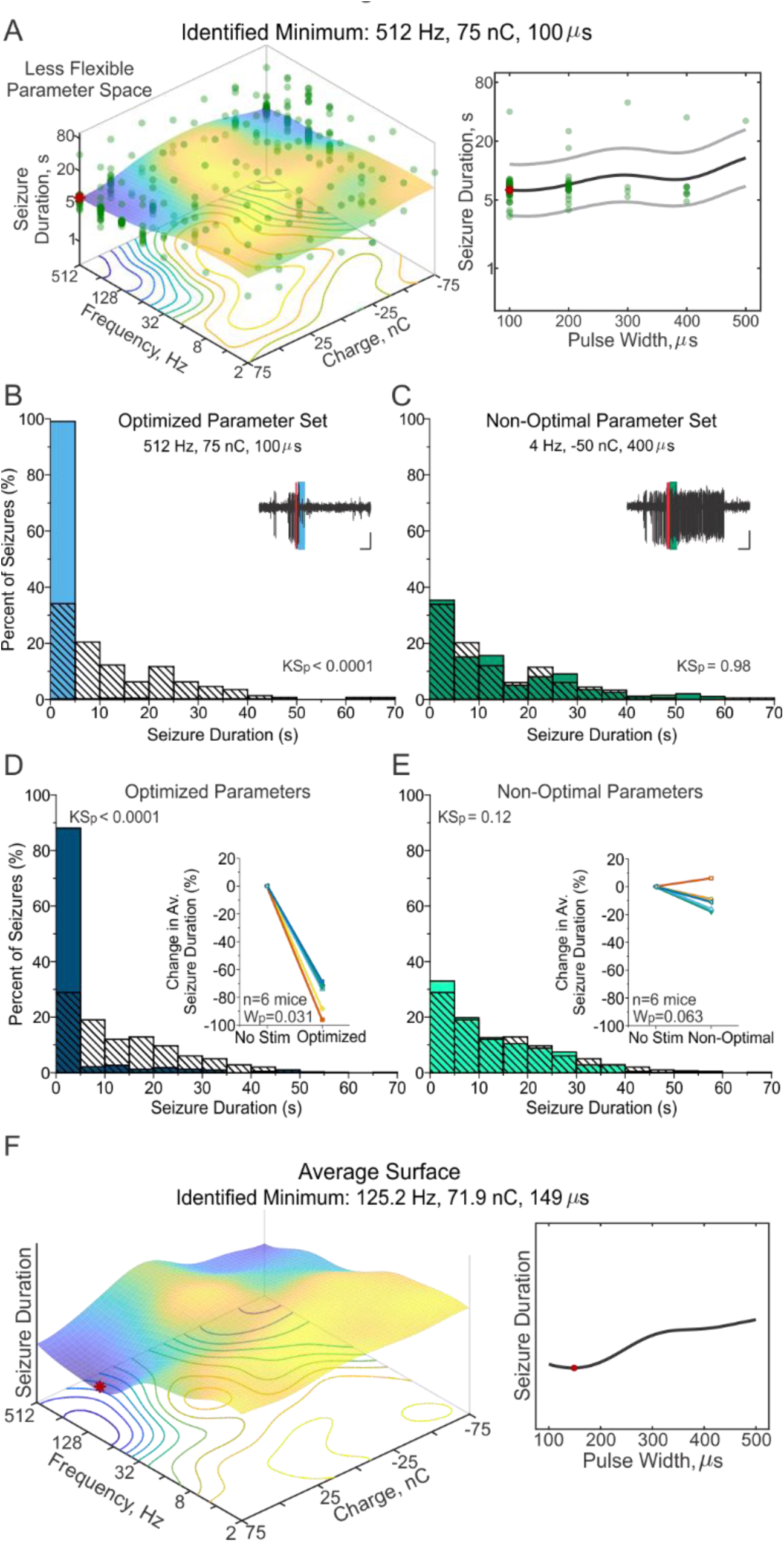
Bayesian optimization identifies effective parameters also under less flexible conditions. (**A**) Example animal’s final response-surface from optimization within the second parameter space, where the number of divisions was reduced from 18 to 9 and 13 to 7 across the frequency and charge dimensions, respectively. Total number of non-zero stimulation combinations was reduced from 1,080 to 270. Additionally, the effective length-scales were increased, causing the surface to be less flexible, or more generalized. (This animal’s more flexible surface optimization can be seen in Supplemental Figure 2). (**B** and **C**) Post-stimulation-onset seizure duration distributions for the example animal shown in (*A*). Light blue bars: events receiving optimized stimulation; dark green bars: events receiving non-optimal stimulation; hashed bars: no-stimulation internal control for comparison. (**B**) A 96% reduction in seizure duration was observed in this animal when the optimized parameter set of 512 Hz, 75 nC, and 100 µs pulse width was applied (p< 0.0001 KS test). Inset: Example seizure event receiving the optimized stimulation (*blue bar*) following detection (*red line*). Scale: 5 s, 0.5 mV. (**C**) This example animal showed a non-significant 6% increase in seizure duration (p = 0.98 KS test) when stimulation was applied with the non-optimal parameter set of 4 Hz, −50 nC, and 400 µs pulse width. Inset: Example seizure event receiving non-optimal stimulation (*green bar*) following detection (*red line*). Scale: 5 s, 0.5 mV. (**D** and **E**) Post-stimulation-onset seizure duration distributions at the group level (100 random seizure events per condition per animal). Dark blue bars: events receiving optimized stimulation; light green bars: events with non-optimal parameters; hashed bars: no stimulation control. (**D**) Seizure duration when stimulation was applied with each animal’s individualized optimized parameter set (*P* < 0.0001 two-sample Kolmogorov-Smirnov test, n=6 mice). Inset: Stimulation with individualized optimized parameters provides consistent, significant, seizure-reduction (78.7±4.4 % reduction in seizure duration, *p* = 0.031 Wilcoxon, significant at the individual animal level in 6 of 6 mice). (**E**) Seizure duration with a non-optimal parameter set unique to each animal (p=0.12 two-sample Kolmogorov-Smirnov test, n=6 mice). Inset: non-optimal parameters do not change seizure duration (9.8±3.5 % reduction in seizure duration, p=0.063 Wilcoxon, not-significant in 5 of 6 mice). Note that each color/symbol represents a different animal, with color coding consistent for panels (*D*) and (*E*). Open symbols: insignificant change at the individual animal level (p>0.05). Closed symbols: significant change for KS and MW test (p<0.01) at the individual animal level. (**F**) Average response-surface, based on the final response-surfaces of the 6 mice. Red dot marks the minimum of the surface at 125.2 Hz, 71.9 nC, and 149 µs pulse width.

These findings indicate that the Gaussian process regression and Bayesian optimization approach taken here was not strongly reliant on the hyperparameters chosen, but rather was sufficiently robust as to find effective stimulation settings with either a more or less flexible surface.

### Effective settings are robust to electrode orientation

Finally, given the striking cytoarchitecture of the cerebellum, in which Purkinje cells’ dendritic trees are oriented parallel to the midline and parallel fibers run orthogonal to the sagittal plane (**Figure 4A**), we additionally tested how the relative orientation of stimulating electrodes could impact outcomes. Our initial testing scheme aligned the electrode feet perpendicular to the midline (**Fig. 4A, left**), such that the prominent path of current would run perpendicular to the parallel fibers (i.e. granule cell axons, which provide excitatory input to Purkinje cells, influencing simple spikes).

**Figure 4.**
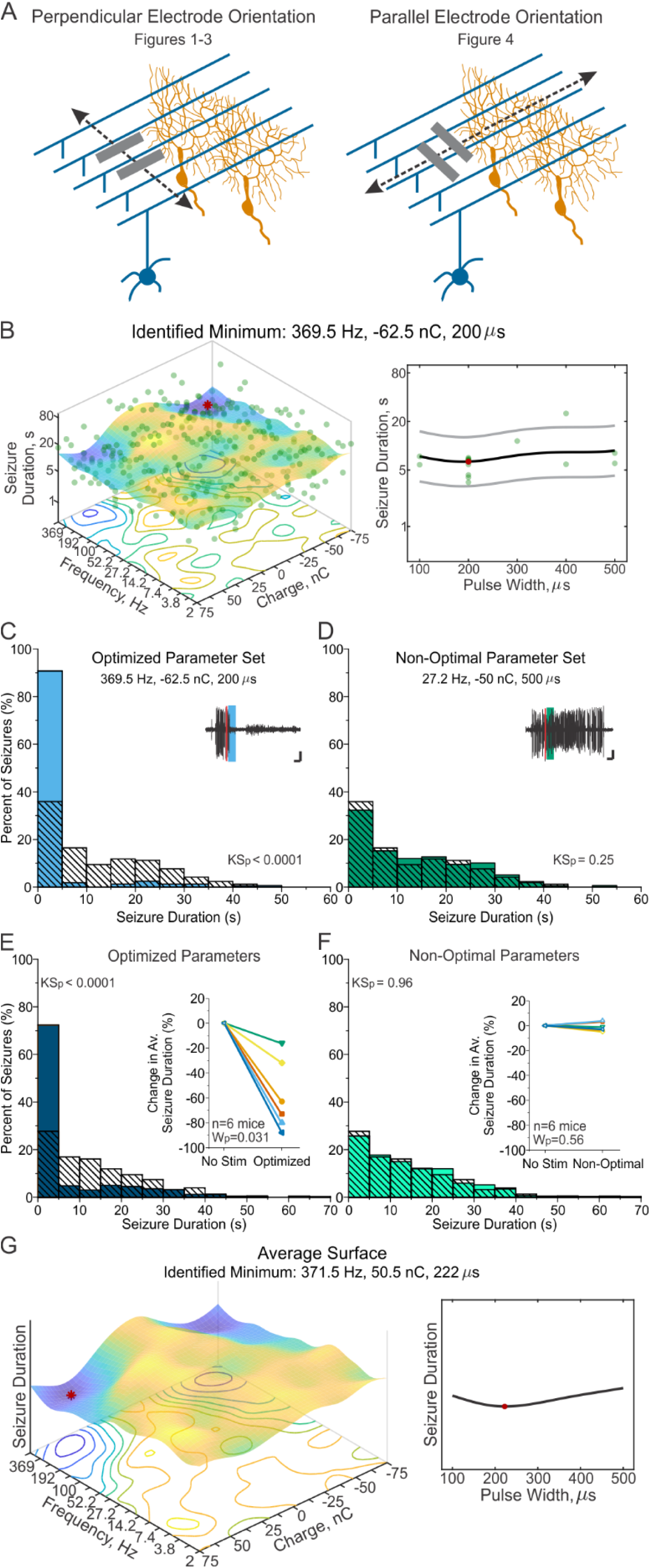
Rotated electrode orientation still permits effective optimization and seizure-control. (**A**) Mice were initially implanted with electrode feet either perpendicular to the midline (*left*). As varying the electrode orientation influences the direction of current flow relative to the stereotyped cytoarchitecture of the cerebellum – where parallel fibers of granule cells (*blue*) run perpendicular to the midline and perpendicular to the nearly-2D dendritic arbors of Purkinje cells (*orange*) – we also tested electrical stimulation of the cerebellar cortex with electrode feet positioned parallel to the midline (*right*). (**B**) Final response-surface from an example animal with a parallel electrode orientation. The minimum of 369.5 Hz, −62.5 nC, and 200 µs pulse width was identified from 1,081 unique possible combinations. (**C** and **D**) Post-stimulation-onset seizure duration distributions for the example animal shown in (*B*). Light blue bars: events receiving optimized stimulation; dark green bars: events receiving non-optimal stimulation; hashed bars: no-stimulation internal control. (**C**) When stimulation was applied with the optimized parameter set of 369.5 Hz, −62.5 nC charge, and 200 µs pulse width, an 80% reduction in seizure duration was observed (p < 0.0001, KS test). Inset: Example seizure event receiving the optimized stimulation (*blue bar*) following detection (*red line*). Scale: 3 s, 0.33 mV. (**D**) Seizure duration when stimulation was applied with the non-optimal parameter set of 4 Hz, −50 nC, and 400 µs pulse width produced a non-significant 4% increase in seizure duration in this animal (p = 0.25 two-sample Kolmogorov-Smirnov test). Inset: Example seizure event receiving non-optimal stimulation (*green bar*) following detection (*red line*). Scale: 3 s, 0.33 mV. (**E** and **F**) Post-stimulation-onset seizure duration distributions across animals (100 random seizure events per condition per animal). Dark blue bars: events receiving optimized stimulation; hashed bars: no-stimulation control; green bars: events receiving non-optimal intervention. Insets: individual animal data. Note that each color/symbol represents a different animal, with color coding consistent for panels (*E*) and (*F*). Open symbols: insignificant change at the individual animal level (p>0.05). Closed symbols: significant change for KS and MW tests (p<0.01) at the individual animal level. (**E**) Seizure duration when stimulation was applied with each animal’s individualized optimized parameter set on average reduced seizure duration by 58.7±11.6% (p=0,031 Wilcoxon, significant in 6 of 6 mice). (**F**) Seizure duration with non-optimal parameter sets did not significantly change seizure duration (average 1±1.5% reduction in seizure duration, p=0.56 Wilcoxon, not significant in 6 of 6 mice). (**G**) Average response-surface, based on the final response-surfaces of 6 mice. Red dot marks the minimum of the surface at 371.5Hz, 50.5 nC, and 222 µs pulse width.

In a second set of animals, we oriented the stimulating electrodes such that the electrode feet were positioned parallel to the midline (**Fig. 4A, right**). In these animals, the main path of current flow would run perpendicular to the dendritic arbors of Purkinje cells (and parallel to parallel fibers). With this rotated orientation of the stimulating electrodes, we again completed Bayesian optimization and Gaussian process regression (using the initial, more flexible, parameter space; **Fig. 4B, Supplemental Figure 6**). Again, this approach allowed identification of predicted effective settings (red dot in example surface in **Fig. 4B**), which in subsequent experiments was confirmed to provide strong seizure inhibition (example animal illustrated in **Fig. 4C**). In contrast, the non-optimal parameter set failed to provide seizure control (illustrated for an example animal in **Fig. 4D**). Across animals, optimized stimulation settings reduced seizure duration by 58.7±11.6% (p=0.031, vs no-stimulation, Wilcoxon; 72% stopping within 5s of intervention, **Fig. 4E, Supplemental Tables 4-5**), while non-optimized stimulation settings produced no seizure inhibition (1±1.5%, p=0.56 vs no stimulation, Wilcoxon; 26% stopping within 5s of intervention; percent seizure reduction non-optimized vs optimized p=0.031, Wilcoxon, **Fig. 4F**). Therefore, as with the perpendicular electrode arrangement, stimulation with a rotated orientation was still able to provide strong seizure inhibition, if the correct combination of stimulation parameters was utilized. Indeed, in a head-to-head comparison between the seizure reduction achieved with optimal settings for the different orientations (Fig.2 data vs Fig. 4 data), there was no significant difference (MW=0.29), suggesting that the orientation of the electrode is not a major determinant of effectiveness of these interventions.

Interestingly, we additionally noted similar surfaces with either electrode orientation scheme (compare **Fig. 4G** to **Fig. 2F**), with fairly high frequencies, fairly strong charges, and somewhat short pulse durations being effective regardless of orientation (**Supplemental Figures 2**,**6; Table 1, Supplemental Table 4**). Averaging across surfaces, the identified minimum (i.e. best combination of settings) for the parallel-to-midline electrode feet orientation had a frequency of 371.5 Hz (compare to 460.1 Hz for our original orientation), a charge of 50.5 nC (compare to 59.7 nC), and a pulse width of 222 µs (compare to 149 µs). The similarities of these optimal settings suggests that not only can seizure inhibition be achieved with either orientation, but also that effective stimulation parameters are similar regardless of the orientation. Therefore, the orientation of electrodes is unlikely to be a critical factor in the success of cerebellar cortex electrical stimulation for seizure inhibition, and therefore likely does not explain variability in past successes when investigating electrical stimulation of the cerebellum for seizure interventions. In contrast, for both orientations, the selection of stimulation settings *was* critical, with some combinations of stimulation settings providing strong seizure inhibition (‘optimized’) and other settings providing no seizure inhibition (‘non-optimal’).

Taken together our results indicate that electrical stimulation of the cerebellum can be an effective approach to seizure inhibition. They further illustrate that effective seizure inhibition is strongly reliant on stimulation settings, providing potential insight into previous mixed findings regarding the effectiveness of cerebellar stimulation for seizure control. Additionally, our work highlights a framework to investigate the importance of stimulation settings, allowing thorough investigation of a large number of possible combinations of settings, in a rationale, data driven, manner.

## Discussion

Previous on-demand optogenetic work suggested strong seizure inhibition via cerebellar-targeted intervention^17-21^, yet previous work targeting the cerebellum via electrical stimulation has suggested extremely mixed results, from no effect ^31,33,40,42,49,53^, to actually worsening the phenotype ^25,29,34,35,39,41,45-47,50,51^, to strong seizure inhibition ^22-30,32,34-39,41,43-45,47,51,52,75^. We hypothesized that electrical stimulation of the cerebellum could provide robust, consistent, seizure inhibition, and that the previous mixed results were likely due to experimental considerations. Specifically, we hypothesized that *how* the intervention was applied would be a critical determinant of successful intervention. We were therefore interested in how stimulation parameters, including stimulation frequency, charge, and pulse width, impacted results. Faced with a very large number of possible combinations of stimulation parameters, we turned to Bayesian optimization^56-58^. Varying frequency from 2Hz to 512Hz with 18 different steps, charge from −75nC to + 75nC with 13 steps, and pulse width from 100µs to 500 µs with 5 steps, resulted in over a thousand different combinations to explore. Such a large number of combinations would have been experimentally intractable to test serially. However, Gaussian process regression and Bayesian optimization allowed a thorough, data-driven exploration of the space, and, importantly, allowed successful identification of parameter settings which were able to provide robust, consistent, seizure inhibition through cerebellar electrical stimulation. This success was not strictly dependent on the fitting conditions, and, surprisingly, was also not dependent on the orientation of the electrodes relative to the cytoarchitecture of the cerebellum. Together, our results highlight the utility of a Bayesian optimization approach to exploring a parameter space, illustrate that the cerebellum can be an effective target for electrical stimulation for seizure inhibition, and ultimately help resolve a decades long controversy, and point of uncertainty, in the field.

Extracting relevant, influential, experimental considerations from past work examining the cerebellum as a potential target for electrical stimulation for seizure inhibition has been remarkably difficult (recently reviewed in Streng and Krook-Magnuson^15^), given factors such as poor reporting of potentially important variables in some studies, and large variability from study to study, with multiple parameters differing. Therefore, a systematic, within-study, rigorous examination of stimulation parameters was critical. By completing our thorough review of a range of frequencies, a range of charge strengths, a range of pulse widths, and, importantly, the various combinations of these, we were able to discern that seizure inhibition required relatively high frequency stimulation and relatively strong charge. Interestingly, the identified minimum (i.e. the “optimal” settings) was often not the highest frequency and highest charge, suggesting that it is not simply a matter of the-more-the-better. While our specific stimulation parameters may not apply to other types of epilepsy, other electrode types or locations, or other species (e.g., humans have much larger brains than mice), it is still interesting to revisit past literature with this knowledge in hand. In doing so, we find that our surfaces and identified optimal settings may explain a large amount of the variability seen in previous studies. For example, high frequency stimulation was effective in inhibiting seizures in studies by Fanardjian^38^ (300Hz), Hutton and colleagues^43^ (200Hz), and Mutani^75^ (100Hz), across a range of seizure models. In contrast, and fitting with the interaction noted in our data between parameters, Grimm and colleagues ^40^ reported that high frequency (250-300Hz) electrical stimulation of the cerebellum failed to inhibit epileptic activity, but, among other differences, used a lower pulse strength as compared to some successful high frequency stimulation studies (i.e. 0.6-0.9V vs e.g. 6-8V in Mutani (1967)^75^). Perhaps most telling, Godlevsky and colleagues^39^ directly compared low (10-12Hz) and high frequency (100-300Hz) stimulation and found that higher frequency stimulation inhibited seizures while low frequency stimulation actually evoked seizures.

In our data set, fitted surfaces tended to predict that low frequency stimulation would result in even worse performance than no stimulation (**Table 1, Supplementary Tables 2, 4**). However, in direct, head-to-head comparisons, we never found a significant worsening of seizure durations with non-optimal stimulation. This suggests that something more may be at play in past studies where electrical stimulation actually made seizures worse ^25,29,34,35,39,41,45-47,50,51^, including the timing of stimulation (our work was entirely on-demand, with stimulation occurring only at the time of seizures, whereas previous studies were done in an open-loop fashion), experimental design (e.g., our work avoided a block design, where natural seizure clustering can make interpretability more challenging), and the potential for plasticity (discussed more below). It is also worth noting that our setup was designed to determine the optimal settings (with increased sampling around potential minima), rather than designed to determine the worst settings; it is possible that even worse settings existed. It should also be noted that it is not surprising that past studies did not do as thorough a review of different parameters and their various potential combinations. Without the regression and optimization approach taken here, examining even half as large of a parameter space would be far too labor intensive. Additionally, the approach taken here of first identifying parameters predicted to be effective, and then performing a secondary direct comparison, allowed us to avoid statistical power pitfalls of thousands of multiple comparisons.

A major advantage of electrical stimulation over optogenetic manipulation is the potential for more immediate translation. As noted before, it seems unlikely that the specific stimulation parameters identified in this study would transfer directly to the human population. However, it is of interest that two human studies report that >50% of patients showed either a reduction in seizures^23^ or full seizure-freedom^26^ when using high frequency stimulation of the cerebellum. It is also important to stress that while extreme caution should be taken when translating the specific settings identified in our study to another population, our results do 1) highlight the potential for robust, consistent, seizure inhibition via cerebellar stimulation, and 2) illustrate that Bayesian optimization can be a successful framework for determining effective stimulation parameters. Such a framework can be applied across settings^59,60,76,77^, including potentially clinical settings, and improvement of interfaces collecting and relaying clinical seizure information is on-going^78^. In our animals, a large number of seizures (over a thousand) were required to adequately map the space and identify minima. While this may seem like far too many to make this a useful approach in a clinical setting, there are a number of important considerations. First, we examined the impact of intervention on non-convulsive, electrographic seizure events. The intrahippocampal KA model, like many human patients^79^, has a high rate of these non-convulsive events^65^. Beyond non-convulsive seizures, human patients will often have a high rate of other epileptiform activity^80^, which can be (and is) targeted in on-demand interventions. Consider, for example, the rate of stimulations (sometimes over a thousand per day) often applied in epilepsy patients with the Neuropace device^5,81^. Similarly, a different surrogate or biomarker for effective stimulation might be available^60^. Moreover, we found that optimization with a more or less flexible surface was able to identify effective settings. The less flexible surface used smaller step sizes and therefore resulted in fewer potential combinations being directly tested. Clinical optimization with a reduced exploration space may allow faster optimization.

Additionally, our results showed similarity in surfaces and identified minima from animal to animal, suggesting that, at least for a similar population and seizure etiology, optimization may not always be required. Indeed, we found no significant differences at the group level whether personalized optimized, group mode, or group average optimal settings were utilized (**Fig. 2G-H**). However, certain animals did appear to show greater sensitivity to the exact stimulation settings used (e.g. animal represented by the light blue triangle in **Fig. 2H**). In the clinical setting, one possibility is that a generic good setting could be used initially, and then, for poor responders, the surface could be explored to identify personalized optimal settings. Even in this circumstance, information from other patients may be able to inform the initial fitting. Our experiments did not make any *a priori* assumptions about the likely surface shape (beyond being multivariate normal); using previously collected data may allow faster convergence. In considering clinical translation, it is also important to note that our study was designed to provide a thorough investigation of the space and avoid over-exploitation of potential minima. In a clinical setting, an approach which focused more on exploitation by maximizing time with effective settings would likely be taken instead (at least after initial fitting).

There are additional considerations that are important to note. First, we optimized only for seizure duration, but other aspects of stimulation, including potential unwanted side effects, can also be very important. For this, in addition to building such considerations directly into the optimization process^59,76^, it may be possible to use unwanted side effects to bound the space examined (akin to e.g. empirically setting limits for DBS for Parkinson’s disease^82^). There are also additional potential variables to optimize across, including total duration of intervention, waveform structure, etc. We have also only examined acute interventions. Over the course of our experiments, non-stationarity did not seem to seem to be a major confound. However, on a longer time scale, plasticity may play a larger role. While providing intervention only in an on-demand manner minimizes stimulation, and therefore potentially plasticity, future work will be needed to examine the impact of chronic cerebellar intervention.

The cerebellum has a striking cytoarchitecture: granule cell axons (i.e. parallel fibers, so called because they run parallel to one another) run orthogonal to and intersect Purkinje cells’ dendritic arbors, which fan parallel to the midline. We tested two different electrode orientations (one perpendicular to the midline and Purkinje cells’ dendrites, one parallel to the midline), and found both could be effective. Moreover, both orientations produced similar parameter surfaces, with similar average optimal parameters. Given the layout of the cerebellum and the impact therefore that direction of current flow can have^83^, we were surprised that electrode orientation did not play a larger role in determining outcomes. Our electrode feet were relatively large (combined exposed surface area of ∼0.35mm^2^), and were separated by a fairly large distance (∼1 mm), and, of course, the current would not have taken a direct linear path between the two feet. Stimulation with a different electrode setup may have produced a greater orientation-specific effect. Additionally, while we examined two different electrode orientations, we examined interventions targeting the cerebellar vermis. Some of the heterogeneity in previous findings may be derived from differences in the location targeted^15^. We found remarkably strong seizure inhibition targeting the midline of the cerebellar cortex (including a 96% decrease in seizure duration in one instance); other regions of the cerebellum may be less effective. Similarly, the parameter surface generated may be dependent on the location. For example, with optogenetic manipulation, excitation or inhibition of the cerebellar cortex (specifically Purkinje cells) is able to inhibit seizures^17^, but at the level of the cerebellar nuclei, only excitation provides seizure control^18,20^. Electrical stimulation of the cerebellar nuclei, instead of the cortex, may result in very different parameter surfaces and optimized settings, with potentially more restricted areas of the parameter surface showing seizure inhibition.

Our work clearly illustrates that electrical stimulation of the cerebellum can be effective, and that the stimulation parameters chosen for intervention are critical for success. Gaussian process regression and Bayesian optimization allowed us to complete a thorough, data-driven, exploration of a very large parameter space, have a visual readout of the process, and successfully identify effective settings. There is nothing that limits the optimization approach taken here to cerebellar-directed interventions, and the problem of many potential combination of settings to choose from is an issue in a large number of contexts^59-64,84^. Even for stimulation targets relatively well established for epilepsy treatment, an optimization approach may allow identification of better stimulation settings, including for non-responders. Beyond potentially missing the best combination of settings, current strategies rely heavily on clinicians modifying stimulation settings in a way that is labor intensive and not as efficiently data driven. An automated data driven approach to determining optimal stimulation settings would reduce the burden on clinicians and likely improve patient outcomes. This is true for epilepsy patients, and for patients receiving or exploring electrical stimulation for a range of neurological disorders and disabilities, including Parkinson’s^61,76^, Essential Tremor^63,85^, chronic pain^86,87^, Obsessive Compulsive Disorder^88,89^, and Depression^90,91^. Our work highlights the potential benefits of a Bayesian optimization approach.

## Abbreviations

DBS: deep brain stimulation
HFOs: high frequency oscillations
KA: kainate
KS: two-sample Kolmogorov-Smirnov statistical test
LFP: local field potential
MW: Mann-Whitney statistical test
RNS: response neurostimulation
VNS: vagal nerve stimulation

## Acknowledgements

The authors would like to thank Dr. Andrew Lamperski for his insight and discussion of Bayesian optimization.

## Supplementary material

Supplementary material is available at *Brain* online.

## References

1. England MJ, Liverman CT, Schultz AM, Strawbridge LM. Epilepsy across the spectrum: promoting health and understanding. A summary of the Institute of Medicine report. Epilepsy Behav. Oct 2012;25(2):266–76. doi:10.1016/j.yebeh.2012.06.016

2. Asadi-Pooya AA, Rostamihosseinkhani M, Farazdaghi M. Seizure and social outcomes in patients with non-surgically treated temporal lobe epilepsy. Epilepsy Behav. Jul 31 2021;122:108227. doi:10.1016/j.yebeh.2021.108227

3. Asadi-Pooya AA, Stewart GR, Abrams DJ, Sharan A. Prevalence and Incidence of Drug-Resistant Mesial Temporal Lobe Epilepsy in the United States. World Neurosurg. Mar 2017;99:662–666. doi:10.1016/j.wneu.2016.12.074

4. Tian N, Boring M, Kobau R, Zack MM, Croft JB. Active Epilepsy and Seizure Control in Adults - United States, 2013 and 2015. MMWR Morb Mortal Wkly Rep. Apr 20 2018;67(15):437–442. doi:10.15585/mmwr.mm6715a1

5. Nair DR, Laxer KD, Weber PB, et al. Nine-year prospective efficacy and safety of brain-responsive neurostimulation for focal epilepsy. Neurology. Sep 1 2020;95(9):e1244–e1256. doi:10.1212/WNL.0000000000010154

6. Ivry RB, Baldo JV. Is the cerebellum involved in learning and cognition? Curr Opin Neurobiol. Apr 1992;2(2):212–6.

7. Strick PL, Dum RP, Fiez JA. Cerebellum and nonmotor function. Annu Rev Neurosci. 2009;32:413–34. doi:10.1146/annurev.neuro.31.060407.125606

8. King M, Hernandez-Castillo CR, Poldrack RA, Ivry RB, Diedrichsen J. Functional boundaries in the human cerebellum revealed by a multi-domain task battery. Nat Neurosci. Aug 2019;22(8):1371–1378. doi:10.1038/s41593-019-0436-x

9. Schmahmann JD. The cerebellum and cognition. Neurosci Lett. Jan 1 2019;688:62–75. doi:10.1016/j.neulet.2018.07.005

10. Argyropoulos GPD, van Dun K, Adamaszek M, et al. The Cerebellar Cognitive Affective/Schmahmann Syndrome: a Task Force Paper. Cerebellum. Feb 2020;19(1):102–125. doi:10.1007/s12311-019-01068-8

11. Zeidler Z, Hoffmann K, Krook-Magnuson E. HippoBellum: acute cerebellar modulation alters hippocampal dynamics and function. Journal of Neuroscience. 2020;(in press)

12. Miterko LN, Baker KB, Beckinghausen J, et al. Consensus Paper: Experimental Neurostimulation of the Cerebellum. Cerebellum. Dec 2019;18(6):1064–1097. doi:10.1007/s12311-019-01041-5

13. Miller JW. The role of mesencephalic and thalamic arousal systems in experimental seizures. Prog Neurobiol. Aug 1992;39(2):155–78.

14. Fountas KN, Kapsalaki E, Hadjigeorgiou G. Cerebellar stimulation in the management of medically intractable epilepsy: a systematic and critical review. Neurosurg Focus. Aug 2010;29(2):E8. doi:10.3171/2010.5.FOCUS10111

15. Streng ML, Krook-Magnuson E. The cerebellum and epilepsy. Epilepsy Behav. Feb 5 2020:106909. doi:10.1016/j.yebeh.2020.106909

16. Kros L, Rooda OH, De Zeeuw CI, Hoebeek FE. Controlling Cerebellar Output to Treat Refractory Epilepsy. Trends Neurosci. Dec 2015;38(12):787–99. doi:10.1016/j.tins.2015.10.002

17. Krook-Magnuson E, Szabo GG, Armstrong C, Oijala M, Soltesz I. Cerebellar Directed Optogenetic Intervention Inhibits Spontaneous Hippocampal Seizures in a Mouse Model of Temporal Lobe Epilepsy. eNeuro. Dec 2014;1(1)doi:10.1523/ENEURO.0005-14.2014

18. Streng ML, Krook-Magnuson E. Excitation, but not inhibition, of the fastigial nucleus provides powerful control over temporal lobe seizures. J Physiol. Jan 2020;598(1):171–187. doi:10.1113/JP278747

19. Streng ML TM, Krook-Magnuson E. Distinct fastigial output channels and their impact on temporal lobe seizures. bioRxiv. 2021;doi: https://doi.org/10.1101/2021.08.18.456836

20. Kros L, Eelkman Rooda OH, Spanke JK, et al. Cerebellar output controls generalized spike-and-wave discharge occurrence. Ann Neurol. Jun 2015;77(6):1027–49. doi:10.1002/ana.24399

21. Eelkman Rooda OHJ, Kros L, Faneyte SJ, et al. Single-pulse stimulation of cerebellar nuclei stops epileptic thalamic activity. Brain Stimul. Jul-Aug 2021;14(4):861–872. doi:10.1016/j.brs.2021.05.002

22. Bidzinski J, Bacia T. [Long-term electrostimulation of the cerebellar cortex in the treatment of epilepsy]. Neurol Neurochir Pol. Jan-Jun 1982;16(1-3):111-4. Chroniczna elektrostymulacja kory modzku w leczeniu padaczki.

23. Chkhenkeli SA, Sramka M, Lortkipanidze GS, et al. Electrophysiological effects and clinical results of direct brain stimulation for intractable epilepsy. Clin Neurol Neurosurg. Sep 2004;106(4):318–29. doi:10.1016/j.clineuro.2004.01.009

24. Cooper IS, Amin I, Gilman S. The effect of chronic cerebellar stimulation upon epilepsy in man. Trans Am Neurol Assoc. 1973;98:192–6.

25. Davis RGE, Engle H., Dusnak A.,. Reduction of Intractable Seizures Using Cerebellar Stimulation. Appl Neuropysiol. 1983;46:57–61.

26. Davis R, Emmonds SE. Cerebellar stimulation for seizure control: 17-year study. Stereotact Funct Neurosurg. 1992;58(1-4):200–8.

27. S. G. Clinical, morphological, biochemical, and physiological effects of cerebellar stimulation. Macel Dekker. 1977;

28. Klun B SV, Strojnik P, Stanic U, Vodovnik L, Zirovnik S. Chronic cerebellar stimulation in the treatment of epilepsy. 1987; Berlin, Heidelberg

29. Levy LF, Auchterlonie WC. Chronic cerebellar stimulation in the treatment of epilepsy. Epilepsia. Jun 1979;20(3):235–45.

30. Sramka M, Fritz G, Galanda M, Nadvornik P. Some observations in treatment stimulation of epilepsy. Acta Neurochir (Wien). 1976;(23 Suppl):257–62.

31. Van Buren JM, Wood JH, Oakley J, Hambrecht F. Preliminary evaluation of cerebellar stimulation by double-blind stimulation and biological criteria in the treatment of epilepsy. J Neurosurg. Mar 1978;48(3):407–16. doi:10.3171/jns.1978.48.3.0407

32. Velasco F, Carrillo-Ruiz JD, Brito F, et al. Double-blind, randomized controlled pilot study of bilateral cerebellar stimulation for treatment of intractable motor seizures. Epilepsia. Jul 2005;46(7):1071–81. doi:10.1111/j.1528-1167.2005.70504.x

33. Wright GD, McLellan DL, Brice JG. A double-blind trial of chronic cerebellar stimulation in twelve patients with severe epilepsy. J Neurol Neurosurg Psychiatry. Aug 1984;47(8):769–74.

34. Babb TL, Mitchell AG, Jr., Crandall PH. Fastigiobulbar and dentatothalamic influences on hippocampal cobalt epilepsy in the cat. Electroencephalogr Clin Neurophysiol. Feb 1974;36(2):141–54.

35. Cooke PM, Snider RS. Some cerebellar influences on electrically-induced cerebral seizures. Epilepsia. Nov 1955;4:19–28.

36. Dauth G DS, Gilman S. Alteration of Purkinje cell activity from transfolial stimulation of the cerebellum in the car. Neurology. 1974;28:654–60.

37. Dow RS, Fernandez-Guardiola A, Manni E. The influence of the cerebellum on experimental epilepsy. Electroencephalogr Clin Neurophysiol. Jun 1962;14:383–98.

38. Fanardjian W DH. An electrophysiological study of cerebellohippocampal relationships in the unrestrained cat. Acta Physiol Acad Sci Hung. 1964;24:321–33.

39. Godlevskii LS, Stepanenko KI, Lobasyuk BA, Sarakhan EV, Bobkova LM. The effects of electrical stimulation of the paleocerebellar cortex on penicillin-induced convulsive activity in rats. Neurosci Behav Physiol. Oct 2004;34(8):797–802.

40. Grimm RJ, Frazee JG, Bell CC, Kawasaki T, Dow RS. Quantitative studies in cobalt model epilepsy: the effect of cerebellar stimulation. Int J Neurol. 1970;7(2):126–40.

41. Hablitz JJ, McSherry JW, Kellaway P. Cortical seizures following cerebellar stimulation in primates. Electroencephalogr Clin Neurophysiol. Apr 1975;38(4):423–6.

42. Hemmy DC, Larson SJ, Sances A, Jr., Millar EA. The effect of cerebellar stimulation on focal seizure activity and spasticity in monkeys. J Neurosurg. May 1977;46(5):648–53. doi:10.3171/jns.1977.46.5.0648

43. Hutton JT, Frost JD Jr., Foster J. the influence of the cerebellum in cat penicillin epilepsy. Epilepsia. Jul 1972;13(3):401–8.

44. Iwata K, Snider RS. Cerebello-hippocampal influences on the electroencephalogram. Electroencephalogr Clin Neurophysiol. Aug 1959;11(3):439–46.

45. A. K. Active arrest mechanisms of epileptic seizures. Epilepsia. 1962;3:329–37.

46. Lockard JS, Ojemann GA, Congdon WC, DuCharme LL. Cerebellar stimulation in alumina-gel monkey model: inverse relationship between clinical seizures and EEG interictal bursts. Epilepsia. Jun 1979;20(3):223–34.

47. Maiti A, Snider RS. Cerebellar control of basal forebrain seizures: amygdala and hippocampus. Epilepsia. Sep 1975;16(3):521–33.

48. Mutani R, Bergamini L, Doriguzzi T. Experimental evidence for the existence of an extrarhinencephalic control of the activity of the cobalt rhinencephalic epileptogenic focus. Part 2. Effects of the paleocerebellar stimulation. Epilepsia. Sep 1969;10(3):351–62.

49. Myers RR, Burchiel KJ, Stockard JJ, Bickford RG. Effects of acute and chronic paleocerebellar stimulation on experimental models of epilepsy in the cat: studies with enflurane, pentylenetetrazol, penicillin, and chloralose. Epilepsia. Jun 1975;16(2):257–67.

50. Reimer GR, Grimm RJ, Dow RS. Effects of cerebellar stimulation on cobalt-induced epilepsy in the cat. Electroencephalogr Clin Neurophysiol. Nov 1967;23(5):456–62.

51. Rucci FS, Giretti ML, La Rocca M. Cerebellum and hyperbaric oxygen. Electroencephalogr Clin Neurophysiol. Oct 1968;25(4):359–71. doi:10.1016/0013-4694(68)90177-6

52. Snider R. Cerebellar modifications of abnormal discharges in cerebral sensory and motor areas. Plenum Press; 1974.

53. Wada JA. Progressive seizure recruitment in subhuman primates and effect of cerebellar stimulation upon developed vs. developing amygdaloid seizures. Bol Estud Med Biol. Oct-1975 Apr 1974;28(8-10):285–301.

54. Ebner TJ, Bantli H, Bloedel JR. Effects of cerebellar stimulation on unitary activity within a chronic epileptic focus in a primate. Electroencephalogr Clin Neurophysiol. Sep 1980;49(5-6):585–99.

55. Sahel JA, Boulanger-Scemama E, Pagot C, et al. Partial recovery of visual function in a blind patient after optogenetic therapy. Nat Med. Jul 2021;27(7):1223–1229. doi:10.1038/s41591-021-01351-4

56. Mockus J. On Bayesian methods for seeking the extremum. presented at: Optimization Techniques IFIP Technical Conference; 1975;

57. Mockus J. Bayesian Approach to Global Optimization: Theory and Applications. Springer Netherlands; 2012.

58. Matheron G. Principals of geostatistics. Econ Geol. 1963;58:1246–1266.

59. Ashmaig O, Connolly M, Gross RE, Mahmoudi B. Bayesian Optimization of Asynchronous Distributed Microelectrode Theta Stimulation and Spatial Memory. Annu Int Conf IEEE Eng Med Biol Soc. Jul 2018;2018:2683–2686. doi:10.1109/EMBC.2018.8512801

60. Park SE, Connolly MJ, Exarchos I, et al. Optimizing neuromodulation based on surrogate neural states for seizure suppression in a rat temporal lobe epilepsy model. J Neural Eng. Jul 16 2020;17(4):046009. doi:10.1088/1741-2552/ab9909

61. Grado LL, Johnson MD, Netoff TI. Bayesian adaptive dual control of deep brain stimulation in a computational model of Parkinson’s disease. PLoS Comput Biol. Dec 2018;14(12):e1006606. doi:10.1371/journal.pcbi.1006606

62. Laferriere S, Bonizzato M, Cote SL, Dancause N, Lajoie G. Hierarchical Bayesian Optimization of Spatiotemporal Neurostimulations for Targeted Motor Outputs. IEEE Trans Neural Syst Rehabil Eng. Jun 2020;28(6):1452–1460. doi:10.1109/TNSRE.2020.2987001

63. Sarikhani P, Miocinovic S, Mahmoudi B. Towards automated patient-specific optimization of deep brain stimulation for movement disorders. Annu Int Conf IEEE Eng Med Biol Soc. Jul 2019;2019:6159–6162. doi:10.1109/EMBC.2019.8857736

64. Smith AC, Shah SA, Hudson AE, et al. A Bayesian statistical analysis of behavioral facilitation associated with deep brain stimulation. J Neurosci Methods. Oct 15 2009;183(2):267–76. doi:10.1016/j.jneumeth.2009.06.028

65. Bouilleret V, Ridoux V, Depaulis A, Marescaux C, Nehlig A, Le Gal La Salle G. Recurrent seizures and hippocampal sclerosis following intrahippocampal kainate injection in adult mice: electroencephalography, histopathology and synaptic reorganization similar to mesial temporal lobe epilepsy. Neuroscience. Mar 1999;89(3):717–29.

66. Bragin A, Engel J Jr., Wilson CL, Vizentin E, Mathern GW. Electrophysiologic analysis of a chronic seizure model after unilateral hippocampal KA injection. Epilepsia. Sep 1999;40(9):1210–21.

67. Armstrong C, Krook-Magnuson E, Oijala M, Soltesz I. Closed-loop optogenetic intervention in mice. Nat Protoc. Aug 2013;8(8):1475–93. doi:10.1038/nprot.2013.080

68. Twele F, Tollner K, Brandt C, Loscher W. Significant effects of sex, strain, and anesthesia in the intrahippocampal kainate mouse model of mesial temporal lobe epilepsy. Epilepsy Behav. Feb 2016;55:47–56. doi:10.1016/j.yebeh.2015.11.027

69. Lisgaras CP, Scharfman HE. Robust chronic convulsive seizures, high frequency oscillations, and human seizure onset patterns in an intrahippocampal kainic acid model in mice. bioRxiv. 2021;doi:10.1101/2021.06.28.450253

70. Hayes MH. Statistical Digital Signal Processing and Modeling. John Wiley & Sons; 1996.

71. Brochu EC, VM; Freitas, N. A tutorial on Bayesian optimization of expensive cots functions, with application to active user modeling and hierarchical reinforcemental learning. 2010;

72. Cogan SF, Ludwig KA, Welle CG, Takmakov P. Tissue damage thresholds during therapeutic electrical stimulation. J Neural Eng. Apr 2016;13(2):021001. doi:10.1088/1741-2560/13/2/021001

73. Bayesian Optimization Algorithm. The MathWorks, Inc. https://www.mathworks.com/help/stats/bayesian-optimization-algorithm.html

74. Bull A. Convergence rates of efficient global optimization algorithms. J Mach Learn Res. 2011;(12):2879-2904.

75. Mutani R. Cobalt experimental hippocampal epilepsy in the cat. Epilepsia. Dec 1967;8(4):223–40.

76. Connolly MJ, Cole ER, Isbaine F, et al. Multi-objective data-driven optimization for improving deep brain stimulation in Parkinson’s disease. J Neural Eng. May 5 2021;18(4)doi:10.1088/1741-2552/abf8ca

77. Connolly MJ, Park SE, Laxpati NG, et al. A framework for designing data-driven optimization systems for neural modulation. J Neural Eng. Dec 3 2020;doi:10.1088/1741-2552/abd048

78. Pal Attia T, Crepeau D, Kremen V, et al. Epilepsy Personal Assistant Device-A Mobile Platform for Brain State, Dense Behavioral and Physiology Tracking and Controlling Adaptive Stimulation. Front Neurol. 2021;12:704170. doi:10.3389/fneur.2021.704170

79. Serrano-Castro PJ, Sanchez-Alvarez JC, Garcia-Gomez T. [Mesial temporal sclerosis (II): clinical features and complementary studies]. Rev Neurol. Apr 1998;26(152):592–7. Esclerosis temporal mesial (II): manifestaciones clinicas y estudios complementarios.

80. Baud MO, Kleen JK, Mirro EA, et al. Multi-day rhythms modulate seizure risk in epilepsy. Nat Commun. Jan 8 2018;9(1):88. doi:10.1038/s41467-017-02577-y

81. Skarpaas TL, Jarosiewicz B, Morrell MJ. Brain-responsive neurostimulation for epilepsy (RNS((R)) System). Epilepsy Res. Jul 2019;153:68–70. doi:10.1016/j.eplepsyres.2019.02.003

82. Butson CR, Cooper SE, Henderson JM, Wolgamuth B, McIntyre CC. Probabilistic analysis of activation volumes generated during deep brain stimulation. Neuroimage. Feb 1 2011;54(3):2096–104. doi:10.1016/j.neuroimage.2010.10.059

83. Ito M. Cerebellar circuitry as a neuronal machine. Prog Neurobiol. Feb-Apr 2006;78(3-5):272–303. doi:10.1016/j.pneurobio.2006.02.006

84. Tervo AE, Metsomaa J, Nieminen JO, Sarvas J, Ilmoniemi RJ. Automated search of stimulation targets with closed-loop transcranial magnetic stimulation. Neuroimage. Oct 15 2020;220:117082. doi:10.1016/j.neuroimage.2020.117082

85. Haddock A, Mitchell KT, Miller A, Ostrem JL, Chizeck HJ, Miocinovic S. Automated Deep Brain Stimulation Programming for Tremor. IEEE Trans Neural Syst Rehabil Eng. Aug 2018;26(8):1618–1625. doi:10.1109/TNSRE.2018.2852222

86. Sheldon B, Staudt MD, Williams L, Harland TA, Pilitsis JG. Spinal cord stimulation programming: a crash course. Neurosurg Rev. Apr 2021;44(2):709–720. doi:10.1007/s10143-020-01299-y

87. Odonkor C, Kwak R, Ting K, Hao D, Collins B, Ahmed S. Fantastic Four: Age, Spinal Cord Stimulator Waveform, Pain Localization and History of Spine Surgery Influence the Odds of Successful Spinal Cord Stimulator Trial. Pain Physician. Jan 2020;23(1):E19–E30.

88. van Westen M, Rietveld E, Bergfeld IO, et al. Optimizing Deep Brain Stimulation Parameters in Obsessive-Compulsive Disorder. Neuromodulation. Feb 2021;24(2):307–315. doi:10.1111/ner.13243

89. Morishita T, Fayad SM, Goodman WK, et al. Surgical neuroanatomy and programming in deep brain stimulation for obsessive compulsive disorder. Neuromodulation. Jun 2014;17(4):312-9; discussion 319. doi:10.1111/ner.12141

90. Dandekar MP, Fenoy AJ, Carvalho AF, Soares JC, Quevedo J. Deep brain stimulation for treatment-resistant depression: an integrative review of preclinical and clinical findings and translational implications. Mol Psychiatry. May 2018;23(5):1094–1112. doi:10.1038/mp.2018.2

91. Roet M, Boonstra J, Sahin E, Mulders AEP, Leentjens AFG, Jahanshahi A. Deep Brain Stimulation for Treatment-Resistant Depression: Towards a More Personalized Treatment Approach. J Clin Med. Aug 24 2020;9(9)doi:10.3390/jcm9092729

