## Supplemental Table for "Optimization of closed-loop electrical stimulation enables robust cerebellar-directed seizure control"

**Supplementary Table 1. Additional summary of data for individual animals presented in Figure 2.**

| | | | Optimized Parameters | | | | Non-Optimal Parameters | | | |
|---|---|---|---|---|---|---|---|---|---|---|
| **Animal ID, Sex**[a] | **# Events for Opt.**[b] | **Time Post-KA (wk)**[c] | **ΔSeizure Duration (%)**[d] | **p-values (KS; MW)**[e] | **ΔTime to Next Seizure (%)**[f] | **p-values (KS; MW)** | **ΔSeizure Duration (%)** | **p-values (KS; MW)** | **ΔTime to Next Seizure (%)** | **p-values (KS; MW)** |
| Yellow ♦ ♀ | 1711 | 9.9 | -82 | 7.9E-47; 8.9E-37 | -28 | 7.7E-05; 0.82 | 5 | 0.46; 0.97 | 35.0 | 0.73; 0.35 |
| D. Orange ■ ♂ | 1996 | 6.4 | -92 | 8.9E-50; 1.3E-43 | 36 | 9.1E-10; 3.5E-04 | -5 | 0.84; 0.91 | 8.5 | 0.98; 0.97 |
| Green ▼ ♂ | 1771 | 6.6 | -87 | 3.0E-73; 5.8E-61 | 11 | 9.2E-08; 0.04 | 4 | 0.43; 0.89 | 30.2 | 0.29; 0.08 |
| L. Orange ● ♂ | 1729 | 17.6 | -71 | 3.6E-36; 1.8E-37 | -29 | 1.8E-08; 0.002 | -12 | 0.13; 0.06 | -2.7 | 0.79; 0.72 |
| D. Blue ◄ ♂ | 1021 | 3.9 | -76 | 1.4E-28; 4.6E-22 | -14 | 2.4E-04; 0.23 | 8 | 0.67; 0.46 | -21.6 | 0.91; 0.47 |
| L. Blue ▲ ♀ | 1334 | 5.0 | -28 | 1.5E-08; 3.1E-06 | -18 | 0.03; 0.04 | -6 | 0.62; 0.54 | -17.2 | 0.72; 0.31 |
| Pink ► ♂ | 1814 | 9.9 | -59 | 3.9E-46; 1.1E-33 | ND[g] | ND | 1 | 0.40; 0.56 | ND | ND |

[a]Animal sex is denoted by ♂ for male, ♀ for female. Color and shape indicate how the animal is presented in Figure 2 (L=light; D=dark)

[b]Number of online events detected and used for Bayesian optimization (opt.) to form the individual final response-surfaces, and identify optimized and non-optimal parameters.

[c]Time between the date of kainic acid (KA) injection and start of on-demand testing corresponding to results presented.

[d]ΔSeizure Duration is calculated relative to the average post-stimulation-onset seizure duration of the no stimulation control. This data and associated statistical analyses are repeated from Table 1 for context.

[e]Significant values ($p$-value < 0.05) are underlined in the table.

[f]ΔTime to Next Seizure is calculated relative to the average time to next seizure for the no stimulation control. A fuller picture of impacts on time to next seizure is better captured by the distributions (see Supplemental Figure 3).

[g]Due to the level of noise in this animal's depth EEG signal, time to next seizure was not analysed (ND=not determined), and this animal did not undergo optimization in the second parameter space.

**Supplemental Table 2. Summary of data for individual animals presented in Figure 3.**

| | Optimized Parameters | | | | | Non-Optimal Parameters | | | | |
|---|---|---|---|---|---|---|---|---|---|---|
| **Animal ID, Sex**[a] | **Freq. (Hz)** | **Charge (nC)** | **Pulse Width (µs)** | **ΔSeizure Duration (%)**[b] | **p-values (KS; MW)**[c] | **Freq. (Hz)** | **Charge (nC)** | **Pulse Width (µs)** | **ΔSeizure Duration (%)** | **p-values (KS; MW)** |
| Yellow ♦ ♀ | 128 | 75 | 200 | -88 | 2.5E-54; 1.1E-44 | 16 | -75 | 500 | -11 | 0.06; 0.07 |
| D. Orange ■ ♂ | 512 | 75 | 100 | -96 | 1.6E-57; 4.6E-45 | 4 | -50 | 400 | 6 | 0.98; 0.79 |
| Green ▼ ♂ | 128 | -75 | 300 | -72 | 3.4E-33; 7.7E-23 | 64 | -75 | 500 | -18 | 0.02; 0.03 |
| L. Orange ● ♂ | 512 | -50 | 500 | -73 | 7.2E-37; 5.6E-39 | 8 | -25 | 400 | -9 | 0.41; 0.29 |
| D. Blue ◀ ♂ | 256 | 75 | 300 | -69 | 3.9E-24; 4.6E-20 | 32 | 25 | 500 | -11 | 0.87; 0.79 |
| L. Blue ▲ ♀ | 256 | 75 | 100 | -74 | 7.3E-37; 3.1E-16 | 256 | 75 | 400 | -16 | 0.11; 0.05 |

[a]Animal sex is denoted by ♂ for male, ♀ for female. Color and shape indicate how the animal is presented in Figure 3 (L=light, D=dark). The same representation is used for each animal across Figures 2 and 3, Tables 1, and Supplementary Tables 2-3.

[b]ΔSeizure Duration is calculated relative to the average post-stimulation-onset seizure duration of the no stimulation control.

[c]Significant values (p-value < 0.05) are underlined in the table.

**Supplemental Table 3. Additional summary of data for individual animals presented in Figure 3.**

| | | | Optimized Parameters | | | | Non-Optimal Parameters | | | |
|---|---|---|---|---|---|---|---|---|---|---|
| **Animal ID, Sex**[a] | **# Events for Opt.**[b] | **Time Post-KA (wk)**[c] | **ΔSeizure Duration (%)**[d] | **p-values (KS; MW)**[e] | **ΔTime to Next Seizure (%)**[f] | **p-values (KS; MW)**[e] | **ΔSeizure Duration (%)** | **p-values (KS; MW)** | **ΔTime to Next Seizure (%)** | **p-values (KS; MW)** |
| Yellow ♦ ♀ | 1185 | 15.7 | -88 | 2.5E-54; 1.1E-44 | -8 | 3.5E-07; 0.24 | -11 | 0.06; 0.07 | 11 | 0.23; 0.28 |
| D. Orange ■ ♂ | 1217 | 9.4 | -96 | 1.6E-57; 4.6E-45 | 104 | 5.2E-29; 4.9E-22 | 6 | 0.98; 0.79 | -27 | 0.38; 0.48 |
| Green ▼ ♂ | 1009 | 12.1 | -72 | 3.4E-33; 7.7E-23 | -25 | 0.042; 0.02 | -18 | 0.02; 0.03 | -6 | 0.35; 0.41 |
| L. Orange ● ♂ | 1664 | 14.6 | -73 | 7.2E-37; 5.6E-39 | -31 | 3.3E-07; 0.0016 | -9 | 0.41; 0.29 | -2 | 0.82; 0.26 |
| D. Blue ◄ ♂ | 1409 | 6.1 | -69 | 3.9E-24; 4.6E-20 | -33 | 2.7E-03; 0.61 | -11 | 0.87; 0.79 | -25 | 0.52; 0.20 |
| L. Blue ▲ ♀ | 1549 | 6.0 | -74 | 7.3E-37; 3.1E-16 | -36 | 4.2E-06; 0.01 | -16 | 0.11; 0.05 | -25 | 0.11; 0.16 |

[a]Animal sex is denoted by ♂ for male, ♀ for female. Color and shape indicate how the animal is presented in Figure 3 (L=light; D=dark).

[b]Number of online events detected and used for Bayesian optimization (opt.) to form the individual final response-surfaces, and identify optimized and non-optimal parameters.

[c]Time between the date of kainic acid (KA) injection and start of on-demand testing corresponding to results presented.

[d]ΔSeizure Duration is calculated relative to the average post-stimulation-onset seizure duration of the no stimulation control. This data and associated statistical analyses are repeated from Supplementary Table 2 for context.

[e]Significant values (p-value < 0.05) are underlined in the table.

[f]ΔTime to Next Seizure is calculated relative to the average time to next seizure for the no stimulation control. A fuller picture of impacts on time to next seizure is better captured by the distributions (see Supplemental Figure 3).

**Supplemental Table 4. Summary of data for individual animals presented in Figure 4.**

| Animal ID, Sex[a] | Optimized Parameters | | | | | Non-Optimal Parameters | | | | |
|---|---|---|---|---|---|---|---|---|---|---|
| | Freq. (Hz) | Charge (nC) | Pulse Width (μs) | ΔSeizure Duration (%)[b] | p-values (KS; MW)[c] | Freq. (Hz) | Charge (nC) | Pulse Width (μs) | ΔSeizure Duration (%) | p-values (KS; MW) |
| L. Blue ▲ ♂ | 369.5 | -62.5 | 200 | -80 | 7.3E-44; 2.5E-34 | 27.2 | -50 | 400 | 4 | 0.25; 0.97 |
| D. Orange ■ ♀ | 369.5 | -75 | 200 | -73 | 1.6E-66; 4.3E-49 | 10.2 | 0 | 100 | 3 | 0.87; 0.53 |
| L. Orange ● ♀ | 266.6 | 75 | 200 | -63 | 4.6E-26; 1.1E-18 | 2.8 | -25 | 400 | -4 | 0.30; 0.86 |
| Yellow ♦ ♂ | 512 | 37.5 | 200 | -32 | 5.7E-09; 1.9E-05 | 37.7 | 25 | 200 | -5 | 0.73; 0.87 |
| Green ▼ ♀ | 138.9 | -37.5 | 200 | -16 | 2.1E-07; 0.0028 | 14.2 | 37.5 | 300 | -1 | 0.57; 0.80 |
| D. Blue ◀ ♂ | 266.7 | 62.5 | 400 | -88 | 1.2E-51; 8.0E-46 | 3.8 | -37.5 | 100 | -3 | 0.65; 0.60 |

[a]Animal sex is denoted by ♂ for male, ♀ for female. Color and shape indicate how the animal is presented in Figure 4 (L=light, D=dark).

[b]ΔSeizure Duration is calculated relative to the average post-stimulation-onset seizure duration of the no stimulation control.

[c]Significant values (p-value < 0.05) are underlined in the table.

**Supplemental Table 5. Additional summary of data for individual animals presented in Figure 4.**

| | | | Optimized Parameters | | | | Non-Optimal Parameters | | | |
|---|---|---|---|---|---|---|---|---|---|---|
| Animal ID, Sex[a] | # Events for Opt.[b] | Time Post-KA (wk)[c] | ΔSeizure Duration (%)[d] | p-values (KS; MW)[e] | ΔTime to Next Seizure (%)[f] | p-values (KS; MW)[e] | ΔSeizure Duration (%) | p-values (KS; MW) | ΔTime to Next Seizure (%) | p-values (KS; MW) |
| L. Blue ▲ ♂ | 1009 | 12.4 | -80 | 7.3E-44; 2.5E-34 | -1 | 4.5E-05; 0.38 | 4 | 0.25; 0.97 | 16 | 0.46; 0.27 |
| D. Orange ■ ♀ | 1217 | 11.6 | -73 | 1.6E-66; 4.3E-49 | -6 | 9.8E-07; 0.24 | 3 | 0.87; 0.53 | 7 | 0.71; 0.94 |
| L. Orange ● ♀ | 1185 | 11.6 | -63 | 4.6E-26; 1.1E-18 | 28 | 1.5E-07; 7.4E-06 | -4 | 0.30; 0.86 | -15 | 1.00; 0.80 |
| Yellow ♦ ♂ | 1664 | 9.9 | -32 | 5.7E-09; 1.9E-05 | -30 | 0.43; 0.48 | -5 | 0.73; 0.87 | 20 | 0.22; 0.82 |
| Green ▼ ♀ | 1409 | 57.3 | -16 | 2.1E-07; 0.0028 | 5 | 0.47; 0.47 | -1 | 0.57; 0.80 | 4 | 0.41; 0.32 |
| D. Blue ◀ ♂ | 1549 | 3.9 | -88 | 1.2E-51; 8.0E-46 | -0.5 | 7.1E-08; 0.18 | -3 | 0.65; 0.60 | 5 | 0.99; 0.75 |

[a]Animal sex is denoted by ♂ for male, ♀ for female. Color and shape indicate how the animal is presented in Figure 4 (L=light; D=dark).

[b]Number of online events detected and used for Bayesian optimization (opt.) to form the individual final response-surfaces, and identify optimized and non-optimal parameters.

[c]Time between the date of kainic acid (KA) injection and start of on-demand testing corresponding to results presented.

[d]ΔSeizure Duration is calculated relative to the average post-stimulation-onset seizure duration of the no stimulation control. This data and associated statistical analyses are repeated from Supplementary Table 4 for context.

[e]Significant values ($p\text{-value} < 0.05$) are underlined in the table.

[f]ΔTime to Next Seizure is calculated relative to the average time to next seizure for the no stimulation control. A fuller picture of impacts on time to next seizure is better captured by the distributions (see Supplemental Figure 3).
