## Supplemental Fig for "Optimization of closed-loop electrical stimulation enables robust cerebellar-directed seizure control"

Supplementary Figure 1

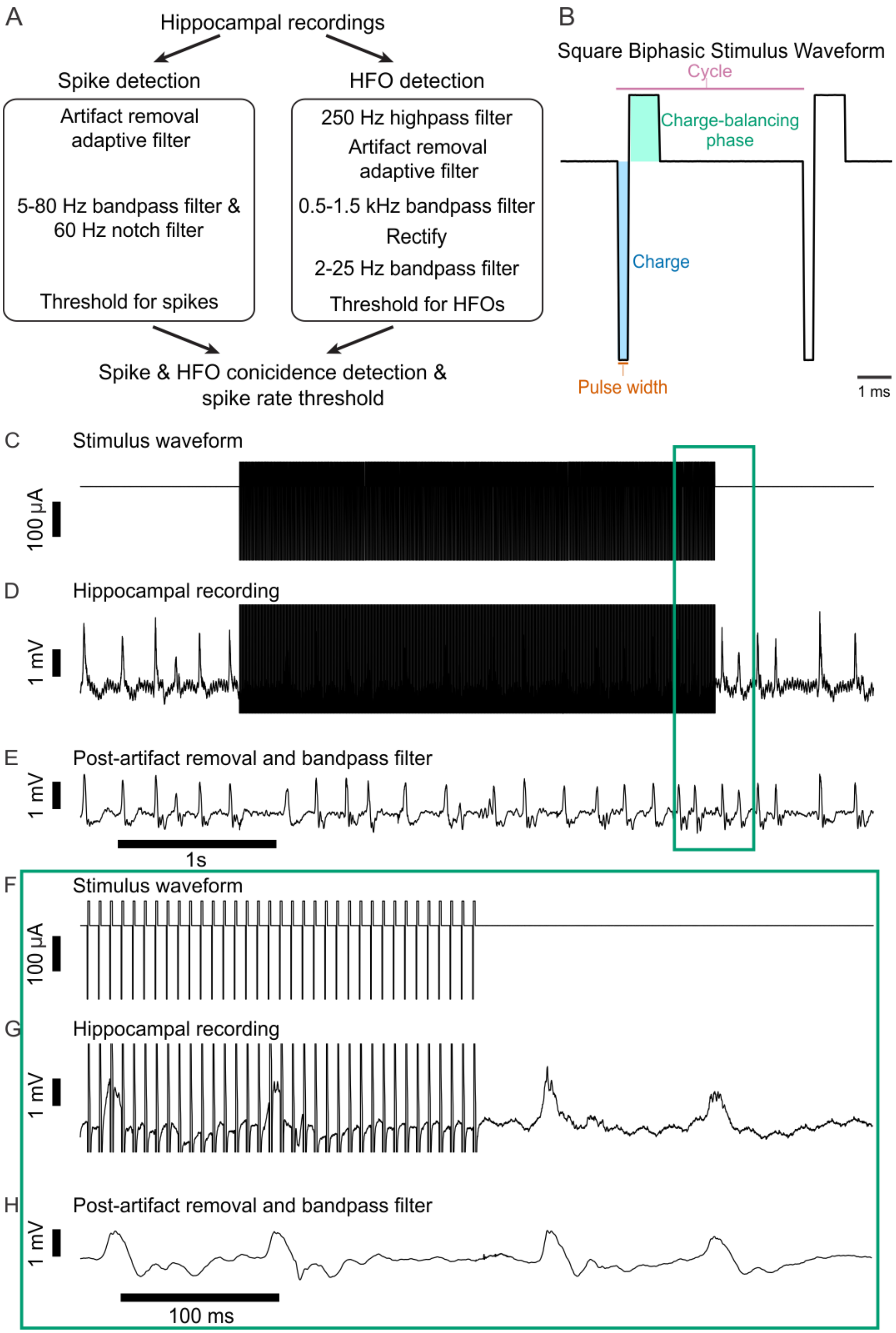

**Supplementary Figure 1. Seizure detection, stimulus waveform, and artifact removal.**

Our closed-loop stimulation and optimization protocol required accurate detection of spikes, even during stimulation. The amplifier gain was set low enough (~100-300x) to ensure the signal would never clip and the sampling rate was sufficiently fast (20,000 samples/s), preserving the superimposed neural and artifact signals. (**A**) We found that each spike was often closely associated with high frequency oscillations (HFOs) (500-1500 Hz). Our detector therefore utilized two parallel pathways to detect both spikes and HFOs. The spike detector began with artifact removal with a least mean squares (LMS) adaptive filter (illustrated further in later panels), followed by bandpass filtering and amplitude thresholding. Several features to isolate spikes, including minimum and maximum inter-spike distance, spike width, and spike amplitude, were set for each mouse, similar to previous online event methods[1]. The HFO detector began with a highpass filter (250 Hz), followed by artifact removal and bandpass filtering (0.5-1.5 kHz). The power envelope of this signal was then captured by rectifying and lowpass filtering the signal. Detection, including whether to require coincident detection of HFOs with spikes, and the minimal spike rate within a two second window for the start and stop of seizures was individualized to allow robust detection with high sensitivity and specificity in each animal. (**B**) Each stimulation pulse consisted of a biphasic waveform, allowing charge balancing. We utilized a 3:1 pulse asymmetry, where the leading phase (*blue*) is followed by a wider charge-balancing phase (*green*) of equal charge, opposite polarity, and three times the pulse width. An example waveform is shown, from a 369.5 Hz frequency, -62.5 nC charge, and 300 µs pulse width stimulus. Note that the charge (nC, *blue*), pulse width (µs, *orange*), and frequency (Hz, reflected in the cycle length, *pink*), were all optimized for seizure-reduction during Bayesian optimization. (**C-H**) An example of on-line artifact removal for spike detection is shown. *F-H* corresponds to the green box in *C-E*, on an expanded time scale. (**C** and **F**) The pulse train control signal generated by the DAC (used to control the stimulator) was additionally recorded by the ADC and used with the adaptive filter. (**D** and **G**) The recorded electrophysiological signals show the stimulation artifact superimposed over the spikes. (**E** and **H**) The LMS adaptive filter uses the stimulus waveform to fit a finite impulse response (FIR) of the artifact. This is similar to fitting a template of the artifact, except that when the waveform is changed, the new template can be calculated automatically using the existing FIR. To improve performance and stability of artifact removal, we trained the LMS adaptive filter two times on each second of data before applying it on the third time to remove the artifact. This allowed us to use a smaller step size for the adaptive filter, improving stability, while rapidly adapting to the artifact, but resulted in a one second delay between seizure detection and onset of stimulation. The artifact removal was very effective, which we attribute to the stimulus and recording (DAC and ADC) being sample-for-sample synchronized with essentially no jitter and therefore the LMS adaptive filter's FIR was well aligned with the artifact. Note that we additionally confirmed that all cases of significant seizure inhibition remained significant if instead of relying on artifact suppression, we assumed any seizure that terminated during stimulation actually terminated at the end of stimulation (i.e., the point where stimulation artifact could no longer obscure ictal spikes). Therefore, our results are robust to any limitations in artifact removal.

Supplemental Figure 2

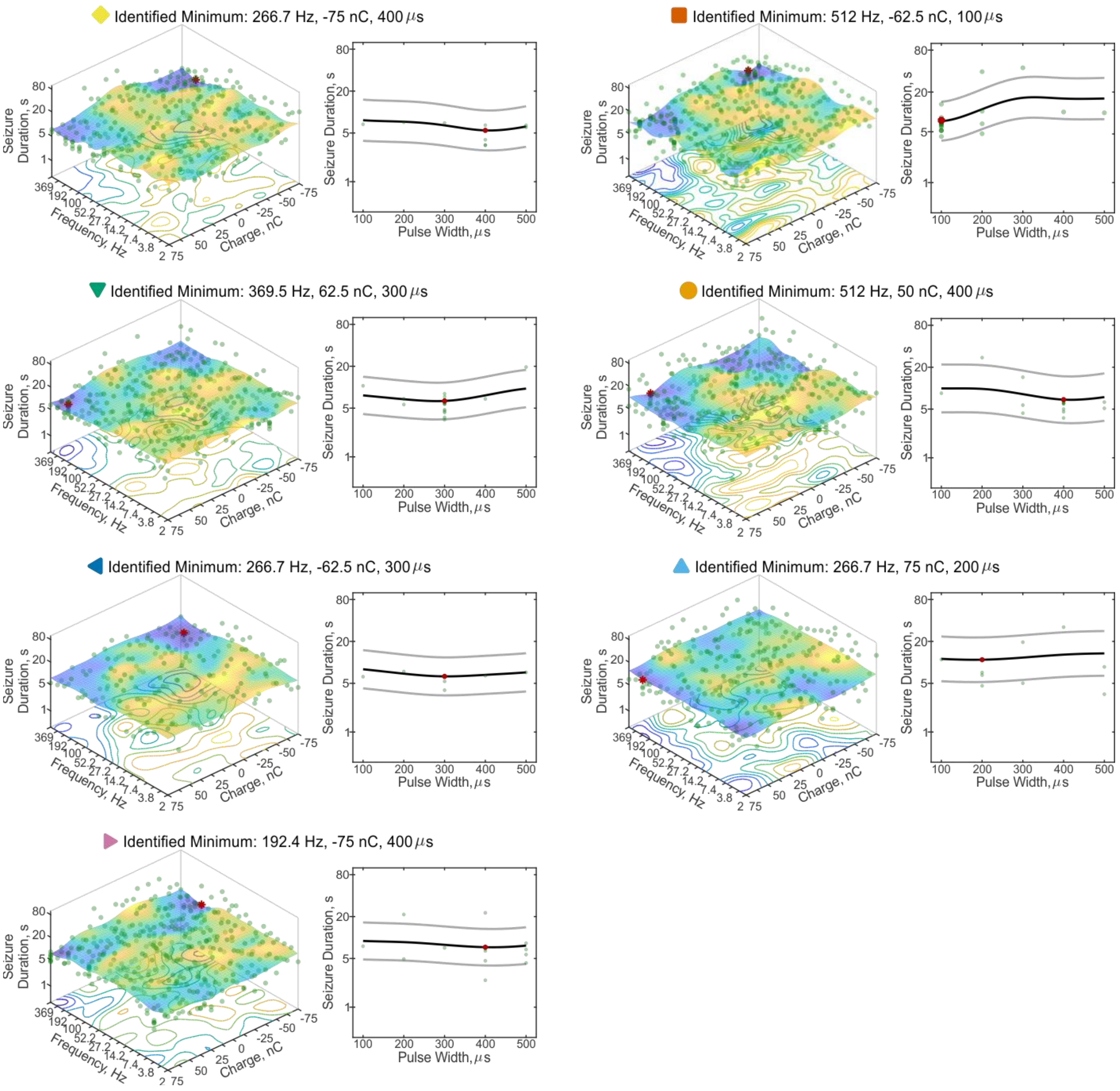


**Supplementary Figure 2. Individual final response-surfaces for each animal presented in Figure 2.** Each final response-surface demonstrates the relative impact of frequency, charge, and pulse width of stimulation on seizure-duration for each animal recorded from with the perpendicular electrode orientation in the original parameter space (n=7 mice). The colored symbol at the top of each plot indicates how the animal is presented in Figure 2, Table 1, and Supplementary Table 1. Light green dots in the plots represent individual recorded events. Cooler colors of the fitted surface in the 3D plots on the left represent shorter seizure durations. The optimized settings for each animal correspond to the minimum of the surface ("Identified Minimum"), marked by the red dot in each plot. For each animal, the 3D plots on the left show the interaction of frequency and charge of stimulation on seizure duration, and the right plot shows the impact of pulse width on seizure duration, at the frequency and charge levels of the minimum.

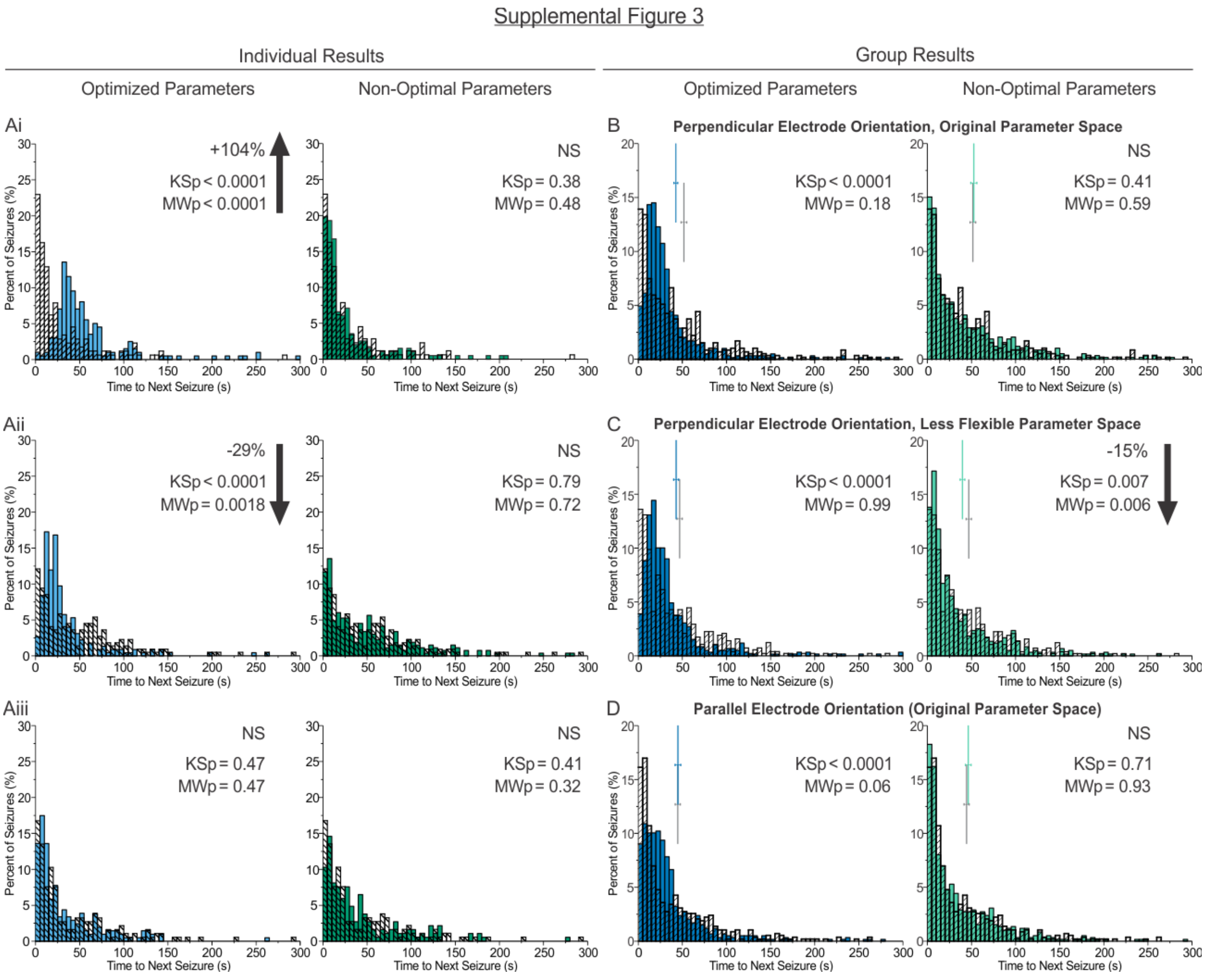


**Supplementary Figure 3. Analysis of time to next seizure.** (**A**) Distributions of time from end of a detected seizure to the start of the next seizure for individual animals. Light blue bars: events receiving optimized stimulation; dark green bars: events receiving non-optimal stimulation; hashed bars: no-stimulation internal control for comparison. Across 18 experiments (n=6 perpendicular electrode orientation, original parameter space; n=6 perpendicular electrode orientation, less flexible parameter space; n=6 parallel electrode orientation, original parameter space), non-optimal stimulation never significantly changed time to next seizure (compared to no stimulation) at the individual level (see Supplementary Tables 1, 3, and 5). Optimal stimulation resulted in a significant change in the average duration of time to next seizure in 9 of 18 instances, with a significant increase in the average time to next (MW<0.05; *Ai, example distribution*) in 4/18, and a significant decrease in the average time to next (*Aii, example distribution*) in 5/18. In seven instances, no change in average duration was noted (MW>0.05), but there was a significant change in the distribution (KS<0.05). In the remaining two instances, there was no change in average duration and no change in the distribution (p>0.05 MW and KS; *Aiii, example distribution*). (**B-D**) Time to next seizure distributions at the group level (100 random seizure events per condition per animal) for experiments illustrated in Figure 2 (**B**), Figure 3 (**C**), or Figure 4 (**D**). Dark blue bars and vertical lines: events receiving optimized stimulation and associated mean ± SEM, respectively; light green bars and vertical lines: events with non-optimal parameters and associated mean ± SEM; hashed bars and gray vertical lines: no stimulation control and associated mean ± SEM.  Note that vertical bars representing means are offset vertically to allow visualization. At the group level, for all experimental groups, optimized stimulation (left side of panels *B-D*) significantly changed the distribution of time to next seizure, but did not change the average time to next seizure. In contrast, non-optimal settings (right panels) typically did not alter the average time to next average times nor distributions, matching the individual animal data, with the exception of group non-optimal stimulation data for the less flexible parameter space (*C*), which resulted in a slight, statistically significant, shift to a decrease in the time to next seizure.

Supplemental Figure 4

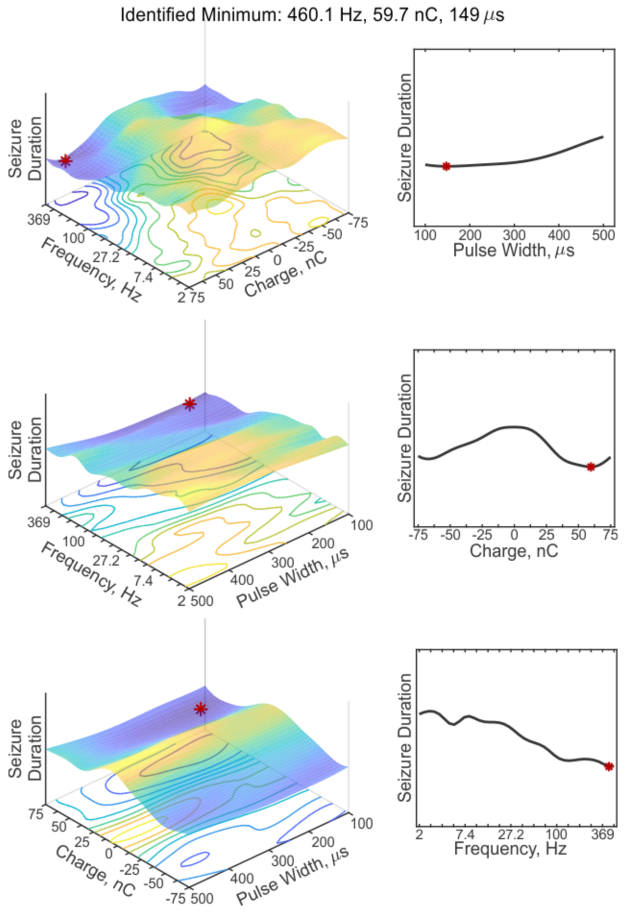


**Supplementary Figure 4. Expanded representation of the averaged final response-surface shown in Fig. 2F.** The final response-surfaces of each animal receiving perpendicular stimulation in the original parameter space (n=7; individual surfaces shown in Supplementary Figure 2) were combined to create a representation of the effects of frequency, charge, and pulse width on seizure duration at the group-level. Due to the high-dimensionality of the data, three planes are shown here. This allows for further examination of the interaction of pulse width with either frequency (*middle row*, *left*) or charge (*bottom row*, *left*), and the impact of charge (*middle row*, *right*) or frequency (*bottom row*, *right*) at the identified minimum (*red dots*) in the other two dimensions.

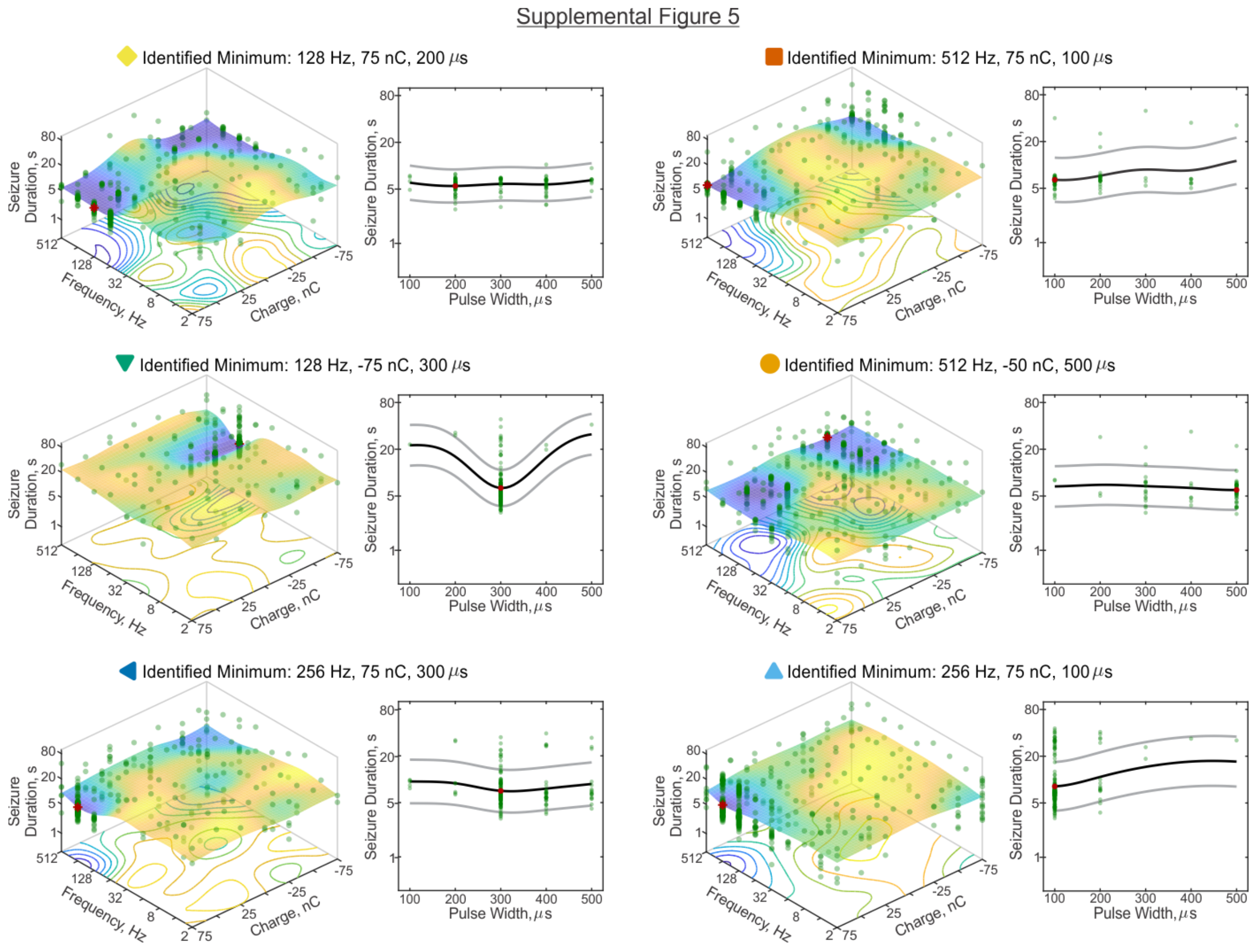


**Supplementary Figure 5. Individual final response-surfaces for each animal presented in Figure 3.** Final response-surfaces for the animals presented in Figure 3, illustrating the results of optimization using a less flexible parameter space. Due to the increased length-scale values and decreased number of tested settings, in the frequency and charge dimensions, the surfaces in this parameter space tend to be smoother (i.e. fewer peaks and valleys; more generalized) than the surfaces in the initial, more flexible, parameter space (compare to surfaces in Supplemental Figure 2). Note that the colored symbol at the top of each plot indicates how the animal is presented in Figure 3, Figure 2, Table 1 and Supplementary Tables 1-3, allowing easy cross-comparison. Note also that the last animal in Supplementary Fig. 2 did not undergo this second optimization due to a poor EEG signal, and is therefore not presented here.

Supplemental Figure 6

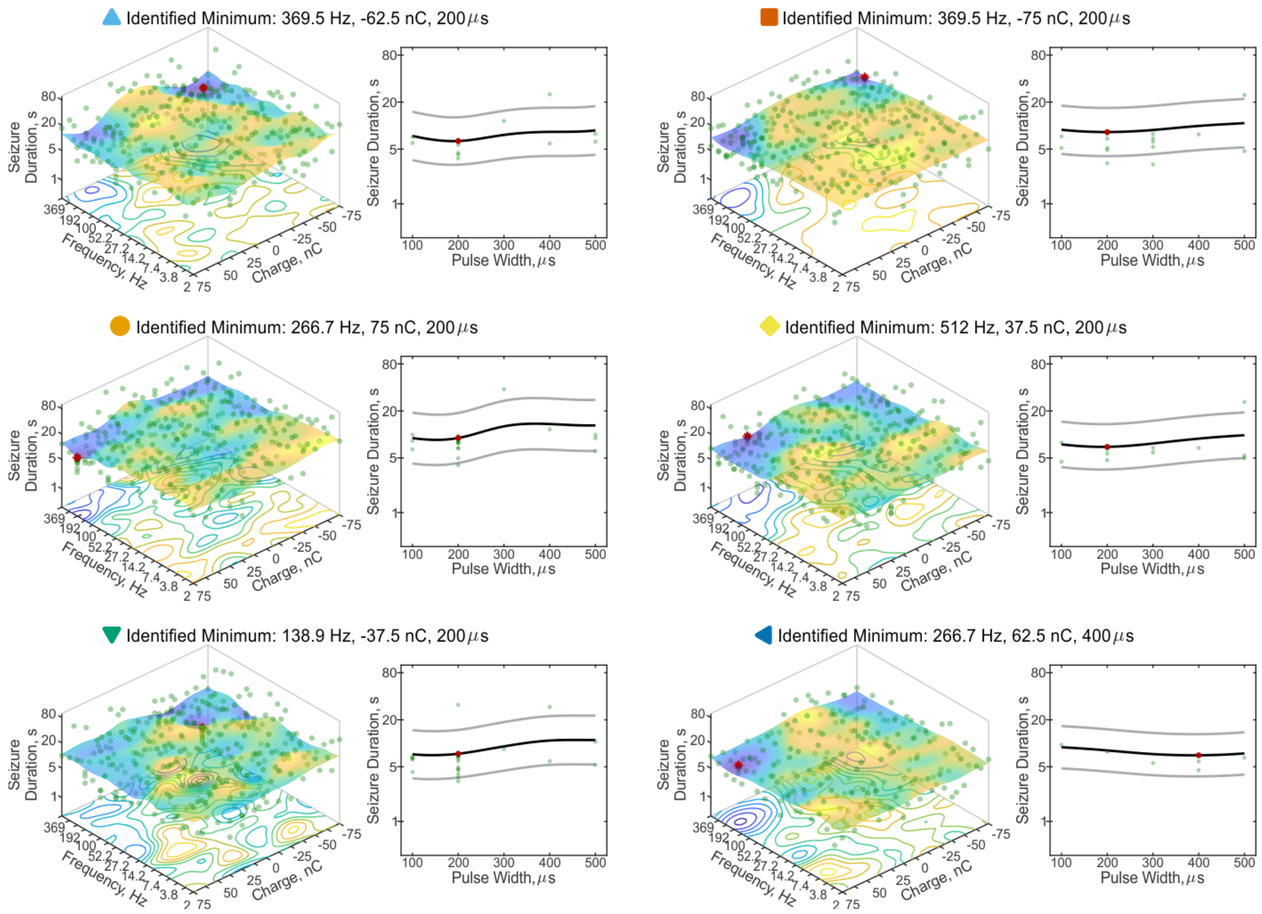


**Supplementary Figure 6. Individual final response-surfaces for each animal presented in Figure 4.** Response-surfaces for animals with the parallel electrode orientation, following optimization in the initial parameter space (same hyperparameters as in Supplementary Fig. 2). The colored symbol at the top of each plot indicates how the animal is presented in Figure 4, and Supplementary Tables 4-5. Note that these are different animals than those presented in Figures 2 and 3 and Supplemental Figures 2-4, despite similar symbols.

**Supplementary Video 1. Response-surface progression for the example animal from Fig. 1.** Using Gaussian process regression, the response surface predicts seizure duration as a function of stimulation parameters. As shown in the video, with each seizure recorded (*green dots*) the surface and the estimated minimum (*red dot*) are updated. The 3D plot on the left shows the interaction of frequency and charge of stimulation on seizure duration at the pulse width of the currently estimated minimum (*cooler colors* indicate predicted shorter seizure durations), and the right plot shows the impact of pulse width on seizure duration, at the frequency and charge levels of the current estimated minimum. After each recorded seizure, Bayesian optimization uses the updated response-surface to determine which parameters to test next to efficiently explore the parameter space.

### References

1. Armstrong C, Krook-Magnuson E, Oijala M, Soltesz I. Closed-loop optogenetic intervention in mice. *Nat Protoc*. Aug 2013;8(8):1475-93. doi:10.1038/nprot.2013.080
